# *SYCP2* translocation-mediated dysregulation and frameshift variants cause human male infertility

**DOI:** 10.1101/641928

**Authors:** Samantha L.P. Schilit, Shreya Menon, Corinna Friedrich, Tammy Kammin, Ellen Wilch, Carrie Hanscom, Sizun Jiang, Sabine Kliesch, Michael E. Talkowski, Frank Tüttelmann, Amy J. MacQueen, Cynthia C. Morton

## Abstract

Infertility is one of the most common disorders for men of reproductive age. To identify novel genetic etiologies, we studied a male with severe oligozoospermia and 46, XY,t(20;22)(q13.3;q11.2). We identified exclusive overexpression of *SYCP2* from the der(20) allele that is hypothesized to result from enhancer adoption. Modeling the dysregulation in budding yeast resulted in disruption of the synaptonemal complex, a common cause of defective spermatogenesis in mammals. Exome sequencing of infertile males revealed three novel heterozygous *SYCP2* frameshift variants in additional subjects with cryptozoospermia and azoospermia. This study provides the first evidence of *SYCP2*-mediated male infertility in humans.

---

Infertility affects 10-15% of couples, making it one of the most common disorders for individuals between the ages of 20-45 years^1^. While many factors contribute to infertility including anatomical defects, gamete integrity, hormonal dysregulation, environmental exposures, age, and certain genetic syndromes, at least 1 in 5 cases of infertility are “unexplained”^2^. Genetic defects may be responsible for many of these idiopathic cases. Indeed, mutations in over 600 genes have been shown to decrease fertility in animal models, and yet few genetic causes of infertility have been validated in humans^3,4^. This results in part from the decreased reproductive fitness of infertile individuals which reduces the number of large families available for genetic analysis in humans as well as the genetic heterogeneity of the disorder^5,6^.

Male factors contribute to about half of all infertility cases, and 40-72% of men lack a specific causal diagnosis^5,7,8^. Given that 30-50% of these cases are estimated to have genetic etiologies, searching for genes involved in unexplained infertility is a rich endeavor^5,7^. Uncovering these novel causes not only informs an understanding of mechanisms regulating fertility, but also provides clinical information to support diagnosis, genetic counseling, and therapeutic intervention.

Currently, the most common genetic testing for male infertility in the clinic involves sequencing of the cystic fibrosis transmembrane conductance regulator *(CFTR)* gene (pathogenic variants found in 80-90% of cases of congenital bilateral absence of the vas deferens), assessing Y chromosome microdeletions (accounting for 5-15% of men with severe oligozoospermia [<5M/ml] or nonobstructive azoospermia), and karyotype analysis (revealing Klinefelter syndrome in 14% of azoospermic men)^3,7,9,10^. In 0.5-1% of severe oligozoospermic or azoospermic men, a karyotype may reveal an apparently balanced reciprocal translocation^11^. While balanced reciprocal translocations may cause subfertility by doubling the risk of miscarriage, the mechanism for how such chromosomal abnormalities may lead to low sperm counts has not been rigorously investigated^12,13^.

We hypothesize that a chromosomal rearrangement may disrupt or dysregulate genes important for fertility in the immediate vicinity of rearrangement breakpoints. By using the well-established Developmental Genome Anatomy Project (DGAP) infrastructure, we initiated a study aimed at identifying new genes important for fertility and exploring an additional explanation for how balanced reciprocal translocations reduce fertility.

## RESULTS

### Clinical report for DGAP230

We sought to identify the genetic etiology of infertility for a male research participant, designated DGAP230, who presented with a two-year history of infertility at age 28. His evaluation showed severe oligozoospermia (<2 M/ml) with normal semen volume. GTG-banded metaphase chromosomes revealed the apparently balanced reciprocal translocation 46, XY,t(20;22)(q13.3;q11.2) (Fig. 1a). Microarray analysis using a 135K-feature whole-genome microarray (SignatureChip Oligo Solution Version 2.0, Signature Genomics Laboratories) indicated no clinically significant abnormalities and no detectable gains or losses of genomic material at either breakpoint. Genetic testing for the Y chromosome microdeletions (YCMD) AZFa, AZFb, and AZFc, as well as *CFTR* pathogenic variants were negative. DGAP230 has no dysmorphic features and has normal serum levels of follicle-stimulating hormone (FSH), luteinizing hormone (LH) and testosterone. Given DGAP230’s normal hormonal levels, normal semen volume and absence of *CFTR* pathogenic variants, we predicted that the etiology of his severe oligozoospermia likely originates in the testes^14^. Based on these initial observations, we hypothesized that the t(20;22) breakpoints disrupt or dysregulate gene(s), which may be important in fertility, perhaps by impacting gametogenesis.

**Fig 1:**
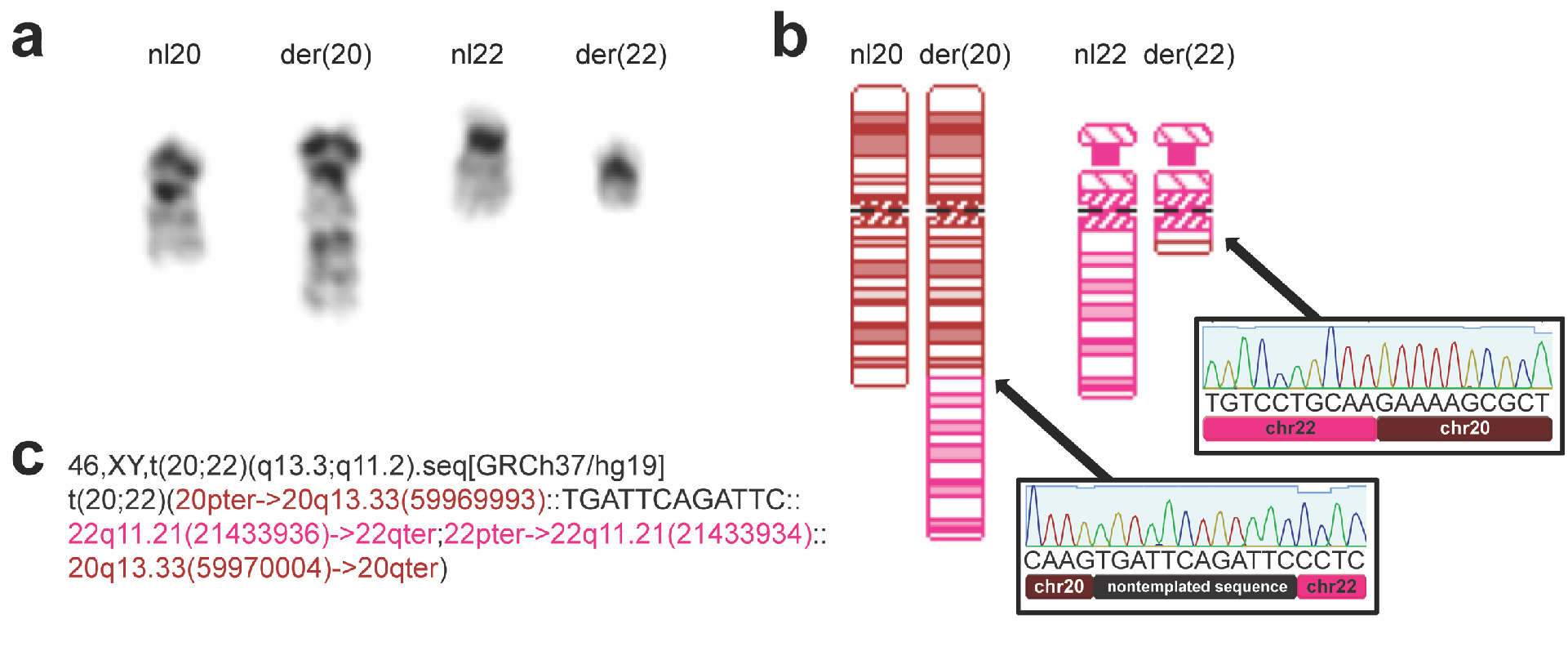
DGAP230 breakpoint characterization. **a**, DGAP230 composite partial karyotype of normal (left) and derivative (right) GTG-banded chromosomes 20 and 22. **b**, The DGAP230 ideogram indicates a simple long arm (q-q) translocation between chromosomes 20 (nl20) and 22 (nl22). Chromatograms from Sanger sequencing show rearrangement breakpoints at nucleotide level resolution. **c**, Next-generation cytogenetic nomenclature is provided for DGAP230^78^. The following software was used to develop this figure: http://www.geneious.com, http://www.cydas.org/OnlineAnalysis/, http://BOSToN.bwh.harvard.edu/.

### Identification of candidate genes for DGAP230’s severe oligozoospermia

To identify gene(s) disrupted or potentially dysregulated by the t(20;22) breakpoints, the translocation was refined to nucleotide resolution using large-insert (“jumping library”) sequencing and subsequent Sanger sequencing (Fig. 1b,c).

Breakpoint locations were then used to identify candidate genes potentially etiologic in DGAP230’s severe oligozoospermia. Genetic regions specifically disrupted by the t(20;22) may be impacted in two ways. If the breakpoint occurs within the open reading frame of any gene, the gene is considered disrupted. Alternatively, positional effects on genes near breakpoints could occur due to loss or gain of regulatory elements in the genetic neighborhood. We defined the potential extent of these effects by assessing the breakpoint-residing topologically associating domains (TADs), compartments of chromatin with frequent interactions in three-dimensional space, because TAD disruption by structural rearrangements can rewire gene expression and induce pathogenicity^15–17^. As a result, our list of candidate genes includes all genes disrupted or potentially dysregulated that reside within the breakpoint-containing TADs.

We computationally assessed predicted TADs by using stem cell Hi-C domains from the Hi-C project (<u>chromosome.dsc.edu/</u>) and converting them to hg19 for comparison to the breakpoints^15^. We identified six genes in the chromosome 20 (chr20) breakpoint-containing TAD and 14 genes in the chromosome 22 (chr22) breakpoint-containing TAD (Fig. 2a). While one gene, *CDH4,* is directly disrupted by the breakpoint on chromosome 20, it is not a strong candidate for severe oligozoospermia. *CDH4* encodes cadherin-4 or R-cadherin, which has predominant expression in the brain and plays a role in retinal axon outgrowth and visual system development^18–20^. The disruption is likely not pathogenic because it has a high haploinsufficiency score (%HI = 37)^21^, suggesting that disruption of only one copy of *CDH4* is not sufficient to induce pathogenicity. In addition, paternally-inherited *CDH4* deletions are reported in DECIPHER, which would be incompatible with a phenotype of male infertility^22^.

**Fig 2:**
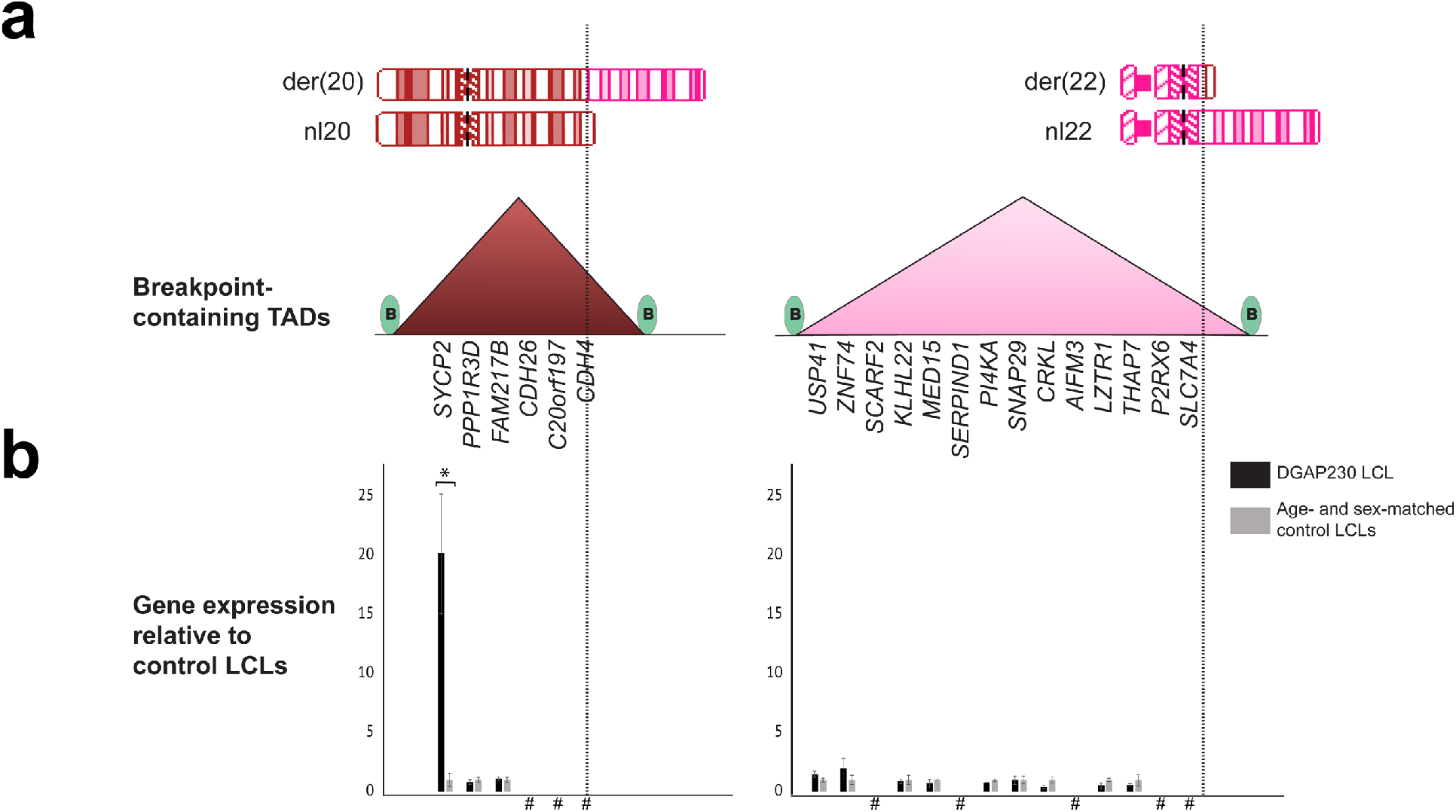
Topologically associating domains disrupted by DGAP230’s t(20;22) define candidate genes. **a**, Vertical lines indicate positions of rearrangement breakpoints across ideograms of normal and derivative chromosomes 20 (mahogany) and 22 (pink). Genes residing in breakpoint-containing topologically associating domains (TADs) are listed below each TAD (triangle) defined by boundary regions (green ovals). **b**, qPCR of candidate genes in the DGAP230 LCL relative to age- and sex-matched control LCLs. Statistical significance was determined by an unpaired twotailed t-test (N=3; p<0.02) and all results display mean ± standard error. # = genes deemed not expressed in LCLs by qPCR standard curve; * = p<0.05.

To determine if DGAP230’s translocation dysregulates candidate genes, we assessed expression of the 20 genes. Using an Epstein-Barr virus-transformed lymphoblastoid cell line (LCL), which has shown prior utility in deducing the effects of structural rearrangements in DGAP subjects^23,24^, we hypothesized that we would be able to detect genes that exhibit constitutive misexpression or overexpression compared to age- and sex-matched controls^20,21^. By assessing expression of every candidate gene by qPCR, we identified one gene on chr20, *SYCP2,* whose expression is increased over 20-fold in DGAP230 LCLs relative to age- and sex-matched controls, a result that was confirmed to be statistically significant by an unpaired two-tailed t-test (p<0.02). No other significant differences were noted in expression between DGAP230 and control LCLs (Fig. 2b).

### Determination of the cytogenetic etiology of *SYCP2* dysregulation

To determine if *SYCP2* misexpression is caused by the t(20;22), we next characterized the mechanism by which *SYCP2* is dysregulated. Given that *SYCP2* is not expressed in karyotypically normal LCLs, we hypothesized that only the der(20)-residing allele would be expressed if the t(20;22) were responsible. To test the hypothesis that only one allele of *SYCP2* is expressed, we identified an exonic polymorphic region in *SYCP2* from DGAP230 genomic DNA and assessed expression of both allelic variants in DGAP230 LCL cDNA^25^. Sequence traces detected only one *SYCP2* haplotype in DGAP230 LCL cDNA, suggesting expression from a single allele in the DGAP230 LCL (Fig. 3a).

**Fig 3:**
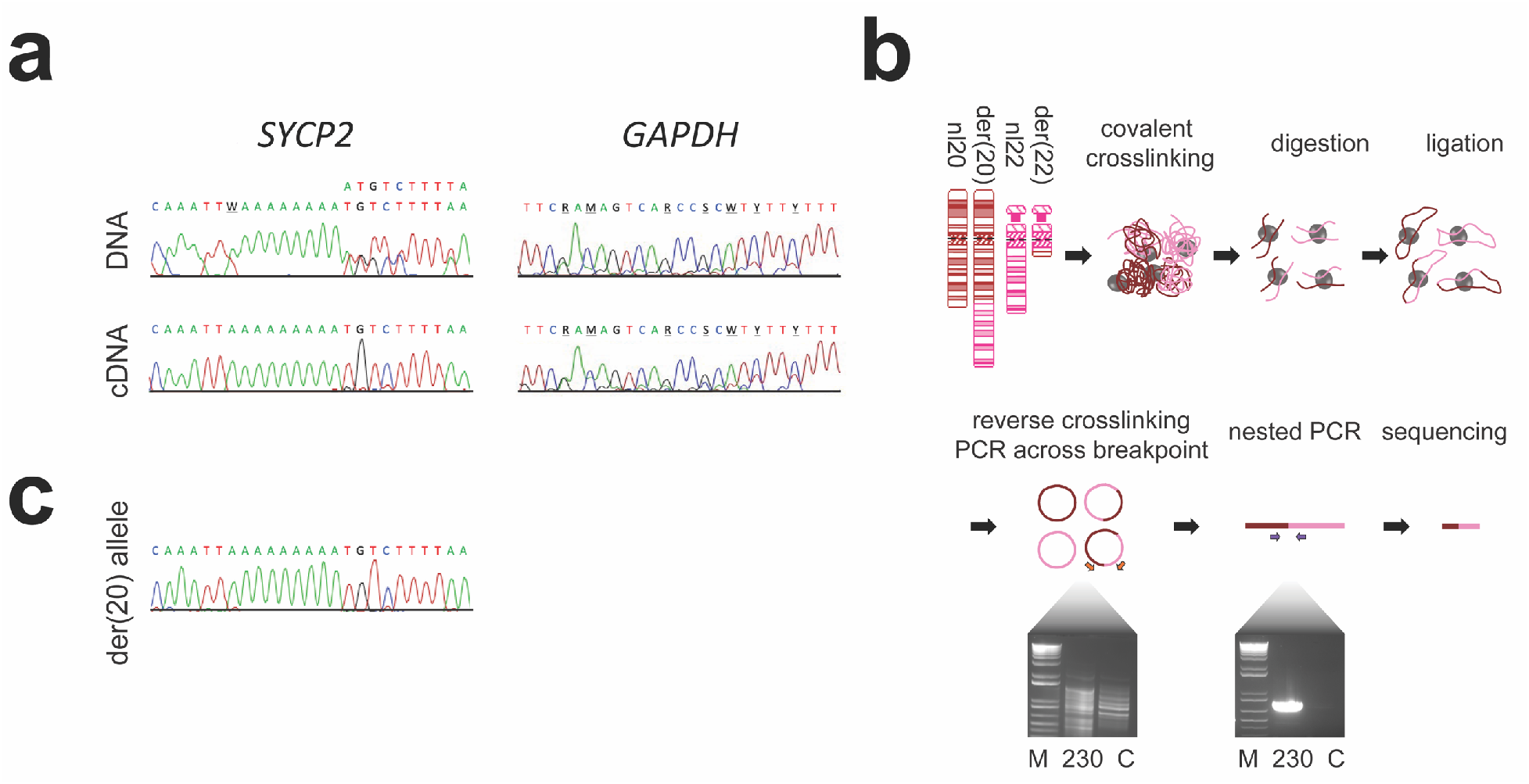
*SYCP2* is overexpressed exclusively from the der(20) allele. **a**, Comparison of DNA and RNA from polymorphic regions in *SYCP2* and *GAPDH* exons. Sequence traces are shown for genomic DNA (top) and cDNA (bottom) from the DGAP230 LCL. The *SYCP2* DNA trace shows heterozygosity at SNPs rs568645874 and rs199662252 while the *SYCP2* cDNA trace only detects one haplotype. In contrast, the *GAPDH* control shows heterogeneity in both DNA and cDNA (SNPs rs45568532, rs551180067 and rs11549332). W = A/T; R = A/G; M = A/C; S = C/G; Y = C/T. **b**, Overview of 3C-PCR, a protocol that couples proximity ligation with breakpoint-spanning PCR to capture *cis* sequences distant from the chromosomal rearrangement^26^. Gel electrophoresis of PCR products from the first PCR across the breakpoint (bottom left) and the second nested PCR using amplification products from the first PCR as a substrate (bottom right) is shown. M = marker; C = control LCLs. **c**, Sanger sequencing traces for the der(20) allele, compared to the DNA and cDNA Sanger sequencing traces in Fig. 3a (N=3).

To examine whether the expressed *SYCP2* allele resides in *cis* with the der(20) translocation breakpoint, we sought to amplify selectively the der(20) allele using primers that span the translocation junction for subsequent Sanger sequencing. To overcome the technical challenge presented by *SYCP2’s* position 1.5 Mb proximal to the translocation breakpoint, we applied 3C-PCR^26^. 3C-PCR capitalizes on principles underlying chromosome conformation capture (3C) to bring fragments containing the translocation junction and the der(20)-residing *SYCP2* allele closer together, thus enabling PCR across the junction of a ligation product including the *cis* allele (Fig. 3b)^27,28^. This resulted in production of amplicons in DGAP230 but not in karyotypically normal LCLs, demonstrating specificity for the t(20;22) (Fig. 3b). Sanger sequencing of this amplicon revealed that it contains the expressed allele (Fig. 3c), suggesting that *SYCP2* expression is detected exclusively from the der(20) allele in the DGAP230 LCL.

Given that misexpression of *SYCP2* derives from the der(20), we hypothesized that *SYCP2* dysregulation is mediated by an “enhancer adoption” mechanism, a long-range *cis*-regulatory mutation that results in a gain of regulatory elements and subsequent promiscuous gene expression^29,30^. To explore this hypothesis, we performed Circular Chromatin Conformation Capture sequencing (4C-seq) using the *SYCP2* promoter as bait in DGAP230 and age- and sex-matched control LCLs and FourCSeq analysis for both normal chromosome 20 (nl20) and der(20) chromosomes^31^. A single statistically significant fragment was identified with 210-fold increased interaction in the DGAP230 LCL (N=3; p_adj_ control = 0.0053), which maps to a genomic region on chr22, 8 Mb downstream of the *SYCP2* promoter (Fig. 4a). A closer look at this region revealed many signatures of enhancer activity including a high ratio of H3K4me1 to H3K4me3, H3K27ac, and two regions that demonstrate DNaseI hypersensitivity in fetal testis tissue (Fig. 4b)^32,33^. Taken together, our findings support an enhancer adoption model, where an active enhancer residing in the segment of chr22 translocated to the der(20) may enter a newly formed chromatin contact encompassing the cis-residing *SYCP2* allele, resulting in illegitimate overexpression.

**Fig 4:**
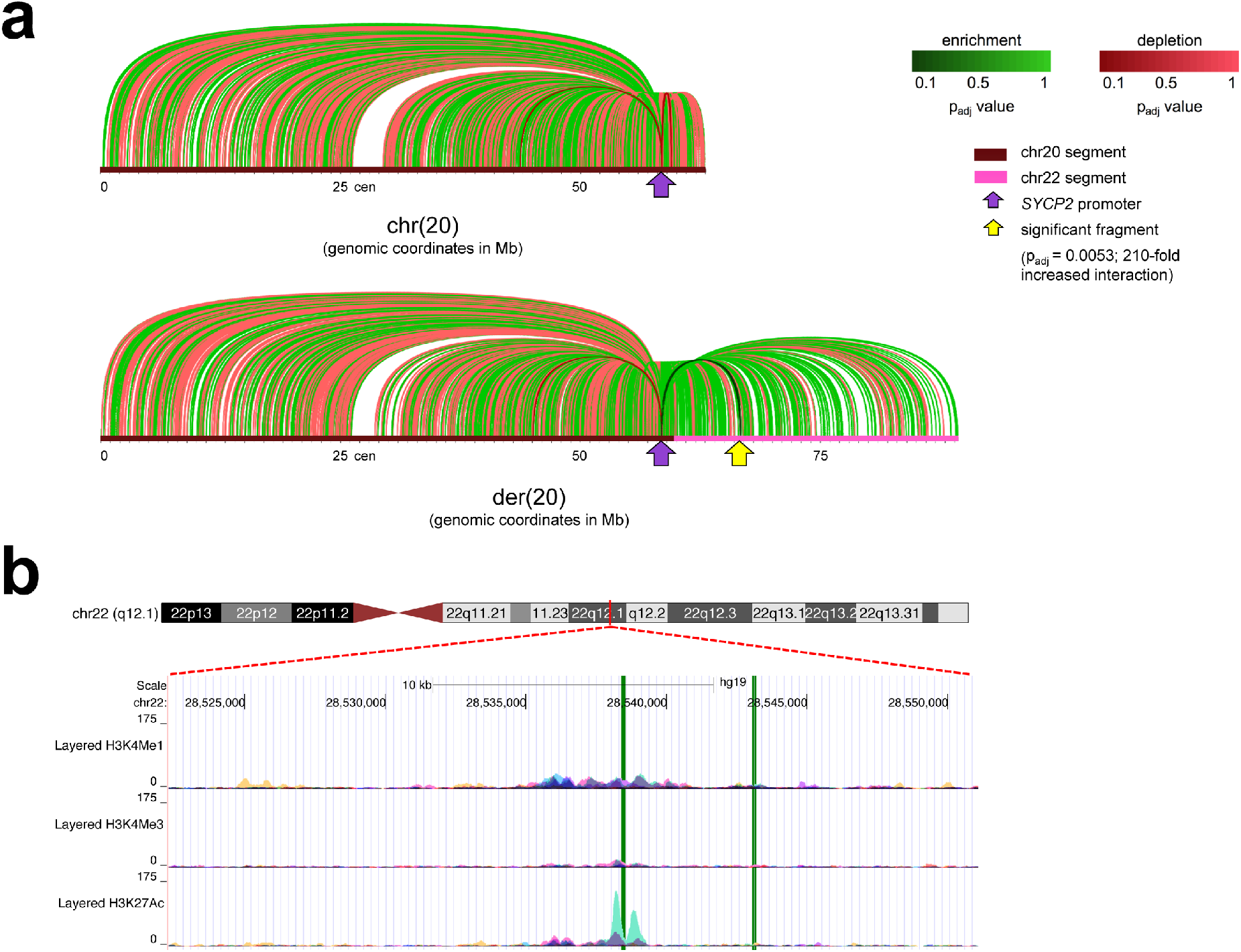
Investigation of chromatin contacts with the *SYCP2* promoter. Circular Chromatin Conformation Capture sequencing (4C-seq) was performed on three biological replicates of the DGAP230 LCL and control LCLs from three different karyotypically normal age- and sex-matched controls. **a**, FourCSeq analysis for chr20 (top) and der(20) (bottom) reveals differential interactions of DNA fragments with the *SYCP2* promoter (purple arrow)^31^. Plots were made in Circos and linearized using Adobe Photoshop. Lines represent each DNA fragment found to interact with the *SYCP2* promoter. A single statistically significant fragment (yellow arrow) with DGAP230 enrichment appears in the der(20) analysis (210-fold enrichment, padj control = 0.0053). cen = centromere. **b**, The significant DNA fragment and flanking enriched regions were consolidated to define a 29 kb putative interacting region. Tracks indicating enhancer activity from seven different cell lines are displayed in the UCSC genome browser and green bars highlight two regions of DNaseI hypersensitivity in fetal testis tissue according to ENCODE^32,33,98^.

In order to implicate the genomic region identified by 4C-seq as the adopted enhancer for *SYCP2* dysregulation, we attempted to delete the putative enhancer region using CRISPR/Cas9 with the goal of assessing the subsequent impact on *SYCP2* expression^34^. Our experiments suggest that the putative enhancer region targeted by CRISPR/Cas9 is important for cell growth or viability (Supplementary Fig. 1), as we were unable to generate DGAP230 LCLs with a deleted putative enhancer region in order to study further its role in *SYCP2* overexpression.

### Analysis of the impact of *SYCP2* misexpression on severe oligozoospermia

We next wanted to understand how *SYCP2* dysregulation can explain DGAP230’s phenotype of severe oligozoospermia. *SYCP2* encodes synaptonemal complex protein 2, a component of the lateral element substructure of the synaptonemal complex (SC)^35^. SYCP2 nonspecifically interacts with the minor groove of DNA and serves as a scaffold for recruiting SYCP3 through its coiled-coil domain, thus facilitating formation of meiotic chromosome axes that are competent to assemble the SC^36–38^. SC assembly (synapsis) is a meiosis-specific process that plays a role in pairing, recombination and segregation of homologous chromosomes during meiosis I^38,39^. SYCP2 is important for spermatogenesis, as male mice homozygous for coiled-coil domain-deficient *Sycp2* exhibit diminished homologous chromosome synapsis, apoptosis within the developing germline, and infertility^36^.

In response to the intriguing role that SYCP2 plays in meiosis and spermatogenesis, we pursued identification of SYCP2 overexpression at the protein level. We found by Western blot that DGAP230 lymphoblastoid cells have over five-times more SYCP2 than age- and sex-matched controls, which achieved statistical significance by an unpaired one-tailed t-test (p<0.0032) (Supplementary Fig. 2).

Based upon DGAP230’s phenotype, his overexpression of *SYCP2* at the RNA and protein levels in LCLs, and current literature on *SYCP2,* we hypothesized that *SYCP2* misexpression may lead to defects in meiosis, resulting in problems with spermatogenesis, thus leading to the phenotype of severe oligozoospermia.

*SYCP2* misexpression may inhibit proper meiosis in DGAP230 because of its role in facilitating homologous chromosome synapsis in meiosis I. It could influence meiosis in two opposing ways. First, it is possible that accumulation of SYCP2 leads to excess synaptonemal complex formation, perhaps preventing proper separation of homologs in anaphase I. Alternatively, excess SYCP2 may lead to an altered stoichiometry that might prevent proper binding of other axial element proteins such as synaptonemal complex protein 3 (SYCP3), causing poor integrity of the synaptonemal complex with resulting asynapsis.

Because meiosis is evolutionarily conserved, the unicellular eukaryote *Saccharomyces cerevisiae* serves as an excellent model organism to study the molecular mechanisms of meiosis^40^. We positioned the yeast functional homolog of *SYCP2, RED1,* under control of the inducible promoter *P_GAL1_*^37^ in a strain background that contains *GAL4-ER,* which induces constitutive strong expression from *P_GAL1_* promoters in the presence of β-estradiol (Supplementary Fig. 3a,b). To learn how induction of excess Red1 impacts its function as an axial element protein, we used immunolocalization to label Red1 on surface-spread meiotic nuclei. While normal Red1 localizes along the lengths of pachytene chromosomes with stretches of staining and nonstaining regions in meiotic prophase nuclei, induced *P_GAL1_-RED1* meiocytes exhibit an aggregation of Red1 protein associated with meiotic chromosomes, which we interpret to be a polycomplex structure (Fig. 5a). We hypothesized that mislocalization of Red1 may influence its ability to serve as a scaffold for the SC. To assess the structural integrity of the SC, we immunostained surface-spread meiotic nuclei for Zip1, a transverse filament in the yeast SC that forms linear structures at the interface of each synapsed chromosome pair (Fig. 5b,c). We discovered diminished Zip1 assembly in *Pgali-RED1* meiotic nuclei compared to control strains, a result that was found to be statistically significant after quantifying the cumulative length of Zip1 per nucleus in a population of 50 cells (Fig. 5d). We confirmed that loss of Zip1 on meiotic chromosomes is not a result of decreased Zip1 expression (Supplementary Fig. 3c). We conclude that overexpression of meiotic chromosome axis/lateral element component Red1 can result in aberrant assembly of synaptonemal complex in budding yeast meiotic nuclei.

**Fig 5:**
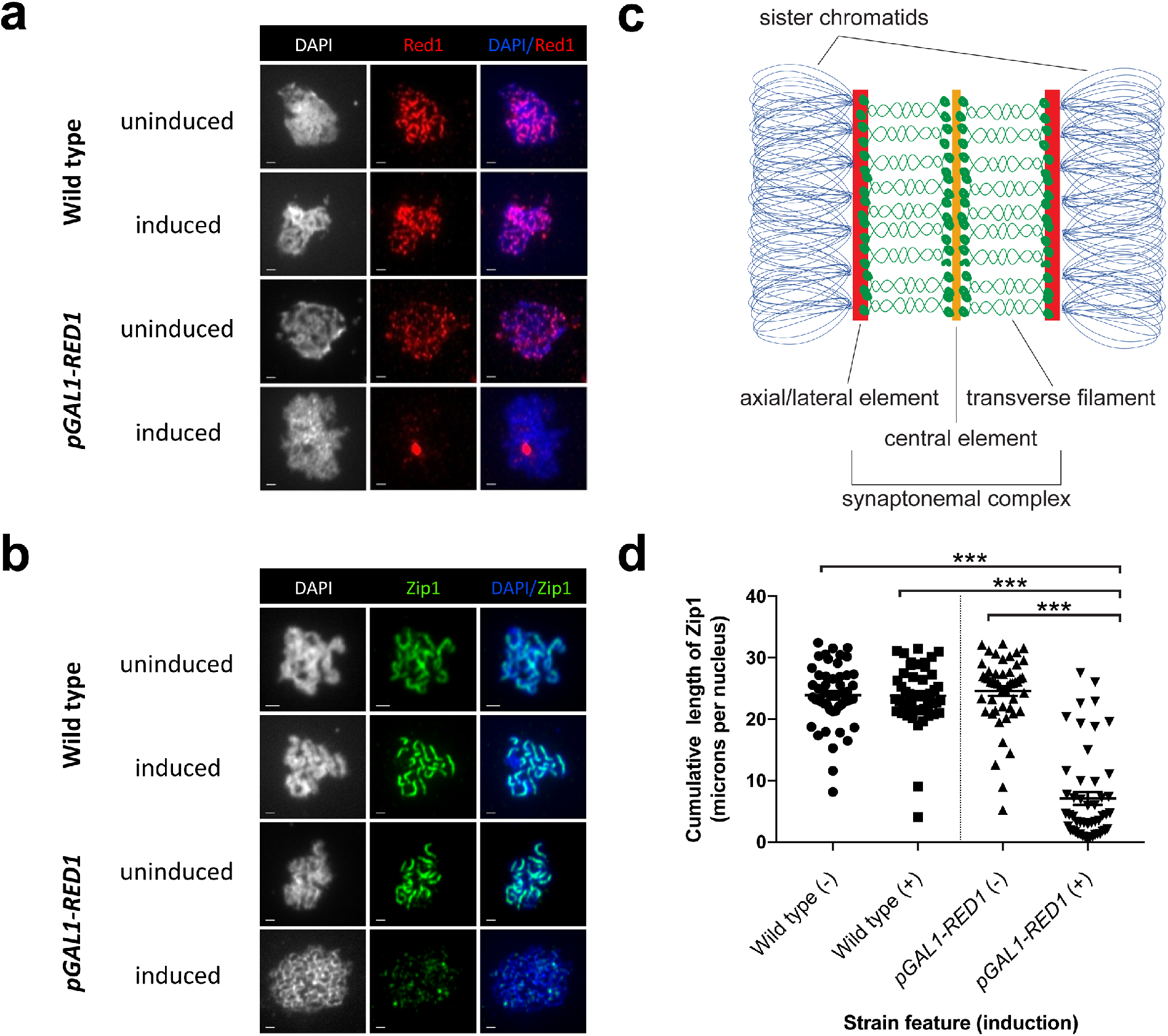
Consequences of Red1 misexpression on synaptonemal complex assembly in S. *cerevisiae*. **a**, Anti-Red1 immuno- and DAPI-staining of surface-spread pachytene chromosomes reveal that constitutive *RED1* induction during sporulation leads to Red1 polycomplex formation (scale bar represents one micron in length). **b**, Anti-Zip1 immuno- and DAPI-staining of surface-spread pachytene chromosomes show that constitutive *RED1* induction during sporulation obliterates Zip1 formation at the interface of homologous chromosomes (scale bar represents one micron in length). **c**, The SC is evolutionary conserved between mammals and budding yeast. SYCP2 and Red1 are axial element functional homologs (red) and SYCP1 and Zip1 are transverse filament homologs (green) in the mammalian and S. *cerevisiae* SC, respectively. The central element substructure is colored yellow. **d**, Quantification of the cumulative length of synaptonemal complex Zip1 in surface-spread meiotic nuclei demonstrates a statistically significant decrease upon induction of *RED1* (Supplementary Fig. 3a,b), as analyzed by Mann-Whitney U test (N = 50). Error bars represent mean ± standard error. *** = p<0.0001.

These results support the possibility that an abnormal abundance of SYCP2 axial element protein disrupts homologous chromosome synapsis in DGAP230, which would be expected to promote germline apoptosis^41^.

### Identification of additional male infertility cases with *SYCP2* loss-of-function variants

Current evidence from our work and the literature suggests that proper SYCP2 dosage is important for its function^36^. In this project, we have found that misexpression of *SYCP2* leads to a loss of function by stoichiometric imbalance. On the other hand, *Sycp2* knockout mice also have an infertility phenotype^36^. Ultimately, it appears that either too much or too little SYCP2 results in a loss of function leading to infertility.

Under the assumption that genetic variants causing infertility would be eliminated from the general population by natural selection, we analyzed allele frequencies of *SYCP2* variants from over 140,000 genomes in the Genome Aggregation Database (gnomAD) browser^42^. We found that *SYCP2* is severely depleted for loss-of-function (LoF) variants including stop-gained and essential splice sites variants, as demonstrated by a *pLI* (probability of loss-of-function intolerance) of 1.000 and *oe* (observed/expected metric) of 0.105^42^. This extreme intolerance to loss of function could be explained by the inability to segregate such variants due to a phenotype of infertility, which would further support *SYCP2* pathogenicity in humans. While we have found that *SYCP2* is significantly depleted for LoF variants, these aggregation datasets lack phenotypic information that would be necessary to assess whether LoF variants lead to infertility. To test the model that LoF *SYCP2* variants are more prevalent in infertile men than in fertile men, we searched for *SYCP2* variants in the exome sequencing (ES) data from the MERGE (Male Reproductive Genomics) study of the Institute of Human Genetics, University of Münster, comprising a male infertility cohort from the Centre of Reproductive Medicine and Andrology (CeRA). This search revealed three males with heterozygous loss-of-function variants at a prevalence of approximately one out of every 200 infertile males.

Participant M1686 presented with a 1.5-year history of infertility at age 29 and was diagnosed with cryptozoospermia (sperm concentration <0.1 M/ml) according to the current World Health Organization (WHO) reference ranges for semen parameters^43^. He had normal testicular volume (right: 17 ml, left: 16 ml [reference range >12 ml]), borderline elevated serum FSH levels (7.1 IU/l [reference range 1-7 IU/l]), and normal LH (9.5 IU/l [reference range 2-10 IU/l]) and testosterone (39.7 nmol/l [reference range >12 nmol/l]). ES and subsequent validation by Sanger sequencing revealed a heterozygous deletion in exon 24 of *SYCP2:* c.2022_2025del causing p.(Lys674AsnfsTer8) (Figure 6a, ClinVar accession no. SUB5364209). This variant is absent in both gnomAD and Trans-Omics for Precision Medicine (TOPMed; https://bravo.sph.umich.edu) databases^42^.

**Fig 6:**
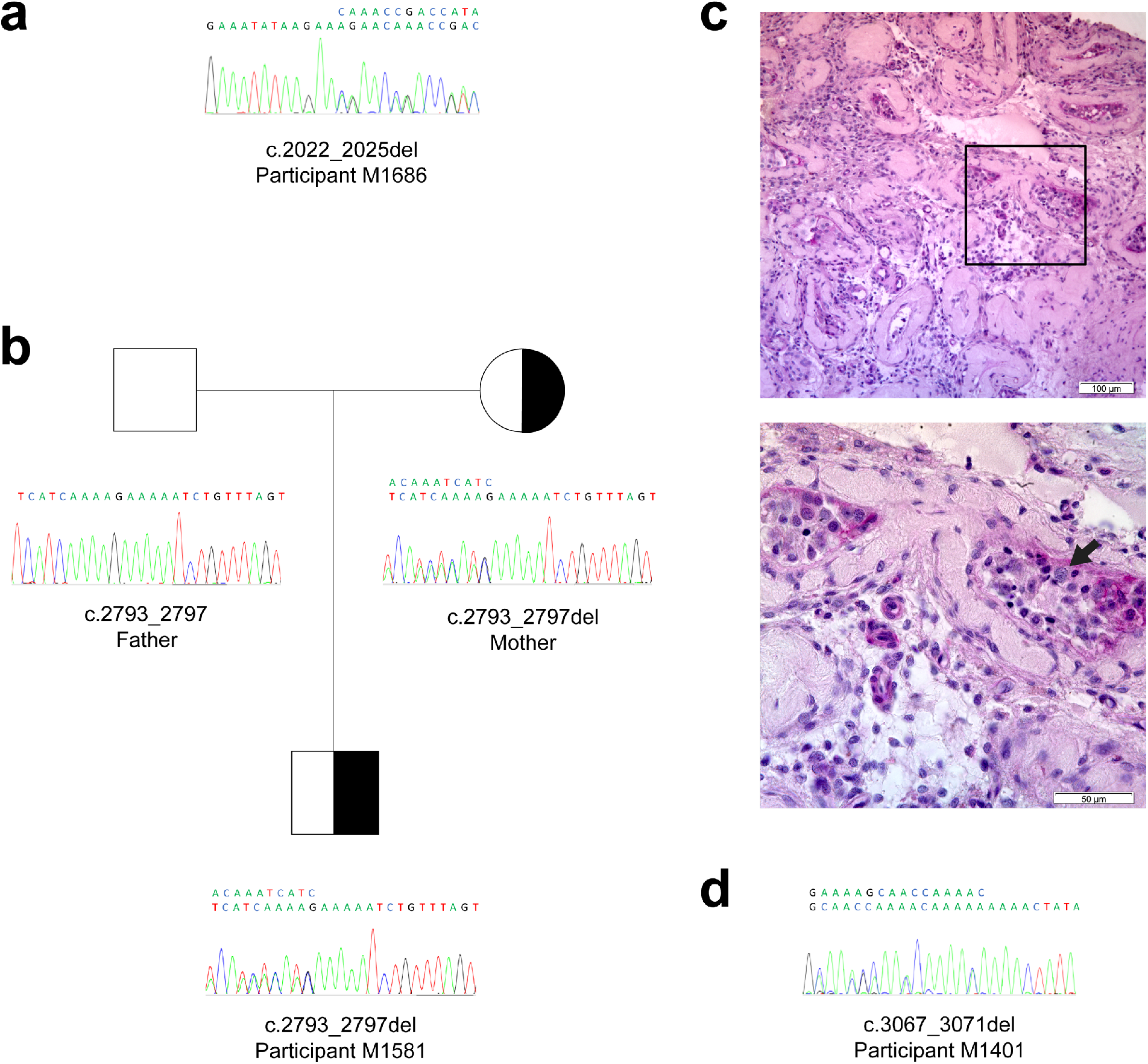
Identification of heterozygous frameshift mutations in *SYCP2* from Münster male infertility cohort participants. **a**, Participant M1686 with cryptozoospermia carries a heterozygous deletion in *SYCP2* (c.2022_2025del, p.(Lys674AsnfsTer8)). **b**, Participant M1581 with cryptozoospermia carries a heterozygous deletion in *SYCP2* (c.2793_2797del, p.(Lys932SerfsTer3)) inherited from his mother. **c**, Histological PAS staining of a testis biopsy from participant M1401 shows a phenotype of meiotic arrest at the pachytene spermatocyte stage leading to nonobstructive azoospermia. The majority of the tubules in an overview section (top) are degenerated to the phenotype of tubular ghosts with a few tubular cross sections that present with spermatocytes. A closer look at the boxed section (magnified in the bottom image) shows disorganized seminiferous epithelium with a single pachytene spermatocyte (black arrow). Scale bar is as indicated. **d**, Participant M1401 carries a heterozygous deletion in *SYCP2* (c.3067_3071del, p.(Lys1023LeufsTer2)).

Participant M1581 from Münster presented with a 2.5-year history of infertility at age 27. He was diagnosed with cryptozoospermia (sperm concentration <0.1 M/ml), had normal testicular volume (right: 14 ml, left: 16 ml), and normal serum levels of FSH (3.3 IU/l), LH (3.0 IU/l) and testosterone (14.5 nmol/l). ES and subsequent validation by Sanger sequencing revealed a heterozygous deletion in exon 31 of *SYCP2:* c.2793_2797del causing p.(Lys932SerfsTer3) (Fig. 6b, ClinVar accession no. SUB5364209). This variant is absent in both gnomAD and TOPMed databases^42^. Segregation was assessed in all available family members, which revealed that the deletion was inherited from the mother while the father is wild type at this position (Fig. 6b).

Participant M1401 presented with a 17-year history of infertility at age 39. He was diagnosed with azoospermia and histopathological analysis from TESE revealed a phenotype of meiotic arrest at the pachytene spermatocyte stage (Fig. 6c). Participant M1401 had borderline testicular atrophy (right: 12 ml, left: 11 ml), elevated serum levels of FSH (44.9 IU/l) and LH (12.4 IU/l), and low serum levels of testosterone (8.2 nmol/l). ES and subsequent validation by Sanger sequencing revealed a heterozygous deletion in exon 33 of *SYCP2:* c.3067_3071del causing p.(Lys1023LeufsTer2) (Fig. 6d, ClinVar accession no. SUB5364209). This variant is absent in both gnomAD and TOPMed databases^42^.

In order to predict the impact of the early termination of *SYCP2* resulting from these variants, we compared them to the coiled-coil domain region identified to be responsible for fertility in the *Sycp2* knockout mouse^36^. By searching for homology to the mouse *Sycp2* coiled-coil domain using a basic local alignment search tool (BLAST; https://blast.ncbi.nlm.nih.gov/), we identified the putative coiled-coil domain of human *SYCP2* in exons 41-44, encoding residues 1408-1505^36,44^. All three termination events reside upstream of the coiled-coil domain, suggesting that these alleles would encode nonfunctional truncated peptides.

## DISCUSSION

Male infertility is a common disorder among reproductive-aged couples and the majority of subjects lack a specific etiologic diagnosis^5^. Understanding the precise causes of male infertility may directly inform therapies for infertile couples. For example, an azoospermic male with a complete deletion of AZFc has a 50% success rate for obtaining sperm by testicular sperm extraction (TESE), while TESE would not be recommended or successful for a man with a complete deletion of AZFa^7^.

Identifying genetic etiologies for human male infertility has been hindered by smaller pedigrees inherent to decreased reproductive fitness and genetic heterogeneity of the disorder. In addition, some genetic evidence of a disorder may not be investigated deeply. For example, balanced reciprocal translocations identified by karyotype in infertile men are rarely followed up beyond reporting a risk for segregation of unbalanced gametes. As a result, a deep investigation into single case studies is critical for uncovering novel genetic etiologies for male infertility.

In this study, we identified a balanced reciprocal translocation in a severe oligozoospermic male designated DGAP230. While it is generally thought that balanced reciprocal translocations may reduce fertility due to production of unbalanced gametes^13^ or meiotic silencing of unsynapsed chromatin^45^, this does not account for the specific phenotype of severe oligozoospermia or azoospermia because the majority of men with balanced reciprocal translocations have normal sperm counts^46^. In addition, men with low sperm counts and a balanced reciprocal translocation have rearrangement breakpoints that sometimes cluster in distinct genomic regions, suggesting that as opposed to a nonspecific mechanism of meiotic segregation, there may be something intrinsic to these genomic regions important for fertility^47,48^.

In the case of DGAP230, the structural rearrangement leads to dysregulation of *SYCP2,* which resides distal to one of the rearrangement breakpoints. This cytogenomic influence on gene expression supports the finding that translocation breakpoints can influence gene expression by dysregulating genes residing within the same TAD^30^. As a result, in men with balanced chromosomal rearrangements and a phenotype of infertility, pathogenesis by specific breakpoints should be considered as an alternative etiology to that of segregation of unbalanced gametes.

While it is known that many different SC proteins have intrinsic ability to self-assemble into polycomplexes when overexpressed, mutated, or expressed in mitotic cells^49–53^, this study demonstrates the first observation of Red1 overexpression leading to its polycomplex formation. We predict that aggregation may be mediated by misexpression before formation of meiotic chromosomes and expression of other axial element proteins as well as the presence of a coiled-coil domain, which facilitates protein complex interactions. Red1 overexpression in budding yeast also decreases double-strand break formation and meiotic recombination initiation, a requirement for SC assembly in budding yeast and most mammals^54^. The resulting asynapsis phenocopies *red1* mutants in *S. cerevisiae* as well as coiled-coil domain-deficient *Sycp2* mice, suggesting that synaptonemal complex formation is sensitive to dosage of axial elements^36,55^.

We believe that our finding of asynapsis resulting from axial element misexpression is directly related to severe oligozoospermia, because asynapsis triggers checkpoint-mediated apoptosis of spermatocytes during spermatogenesis^41,56,57^, which reduces sperm count and has been shown to lead to male-specific infertility^36^. Therefore, DGAP230’s phenotype of severe oligozoospermia and infertility is likely due to asynapsis-triggered cell death in spermatocytes.

The identification of three novel frameshift variants in *SYCP2* from men with cryptozoospermia and meiotic arrest further supports the role of *SYCP2* in human male fertility. All variants are extremely rare consistent with the inability to segregate these mutations in the general population due to a phenotype of infertility. This is also supported by the maternal inheritance of the variant in participant M1581, as *SYCP2*-mediated pathogenicity has been shown to cause male infertility but not female infertility in a mouse model^36^.

Stop codons resulting from the three frameshift variants reside upstream of the coiled-coil domain, which is critical for functionality of the protein^36^. It is important to note, however, that all cases represent heterozygous LoF variants which would support an autosomal dominant disease model. This is discordant with the *SYCP2* knockout mouse model that only demonstrates a male infertility phenotype in homozygotes^36^. However, haploinsufficiency discordance has been observed between human and mouse for the male infertility gene *SYCP3,* the other mammalian axial element in the synaptonemal complex as well as numerous other mouse models^58,59^. *SYCP2’s pLI* > 0.9 and *oe* <0.35 also support a haploinsufficient model, as extreme intolerance to LoF according to constraint analysis is strongly correlated with haploinsufficiency^42^. While less common than autosomal recessive forms of male infertility, autosomal dominant forms have been identified for pathogenic variants in *HIWI^60^, KLHL10^61^ PLK4^62^, SYCP3^58^, SPINK2^63^, NR5A1^64^,* and *DMRT1*^65^.

Another potential concern is that the phenotypes of severe oligozoospermia, cryptozoospermia and meiotic arrest are distinctive from each other. It is possible that the differences reflect variable expressivity, as has been observed in the male infertility genes *DBY* (Sertoli cell-only syndrome and severe hypospermatogenesis), *KLHL10* (severe oligozoospermia and oligozoospermia), *TAF4B* (azoospermia and oligozoospermia), *TDRD9* (azoospermia and cryptozoospermia), and *TEX11* (complete meiotic arrest and mixed testicular atrophy)^61,66–70^.

It is well known that homologous chromosome synapsis is critical for spermatogenesis. Indeed, several genes implicated in human male infertility encode proteins that are members of the synaptonemal complex *(SYCP3* and *SYCE1)* or are otherwise required for synapsis *(SPO11, MEIOB, TEX11,* and TEX15)^6,58,67,70–73^. Before this study, *SYCP2* was considered a strong candidate gene for human male infertility because it encodes a protein that interacts directly with protein products of the human male infertility genes *SYCP3* and *TEX11,* serves as an axial element in the synaptonemal complex, and is required for male fertility in the mouse^36,37,74^. DGAP230 and the three participants from the Münster cohort represent the first cases of putative *SYCP2*-mediated male infertility in humans.

## ONLINE METHODS

### DGAP participant recruitment

Subjects DGAP230, with 46,XY,t(20;22)(q13.3;q11.2), and DGAP278-02, a karyotypically normal age- and sex-matched control, were enrolled through the Developmental Genome Anatomy Project (DGAP). DGAP obtained informed consent, medical records and blood samples under a protocol approved by the Partners HealthCare Systems Institutional Review Board.

### Acquisition of lymphoblastoid cell lines

Epstein-Barr virus-transformed lymphoblastoid cell lines (LCLs) were generated at the Genomics and Technology Core in the Center for Human Genetic Research at Massachusetts General Hospital (Boston, MA, USA). Confirmatory GTG-banded karyotyping was performed on the DGAP230 lymphoblastoid cell line at the Brigham and Women’s Hospital CytoGenomics Core Laboratory (Boston, MA, USA). Two additional karyotypically normal age- and sex-matched control LCLs, GM20184 and GM20188, were obtained from the National Institute of General Medical Sciences (NIGMS) Human Genetic Cell Repository at the Coriell Institute for Medical Research (Camden, NJ, USA).

### Genome jumping library sequencing and analysis

Large-insert (“jumping library”) genome sequencing and subsequent Sanger sequencing identified the precise breakpoints of the DGAP230 chromosomal rearrangement (using primers SS_58F-SS_61R from Supplementary Table 2)^75–77^. Chromatograms were analyzed in Geneious (Version 7.0, Biomatters) and described using next-generation cytogenetic nomenclature^78^.

### Delineation of topologically associating domains disrupted by DGAP230’s rearrangement breakpoints

Topologically associating domains (TADs) disrupted by the breakpoints in DGAP230 were identified according to human embryonic stem cell Hi-C domains from the Hi-C project^15^. The University of California Santa Cruz Genome Browser was used to delineate genes residing in these regions^79^.

### Culturing of lymphoblastoid cell lines

Lymphoblastoid cells were grown at 37°C, 5% CO2, in R10 media (Roswell Park Memorial Institute [RPMI] 1640 Medium without L-Glutamine supplemented with 10% heat-inactivated fetal bovine serum (FBS), 100 U/ml Penicillin-Streptomycin-Glutamine, and 2 mM L-glutamine [all Gibco]). Cells were fed fresh R10 every two or three days. For clonal expansion of single lymphoblastoid cells in the CRISPR/Cas9-mediated genome editing experiment, 30% 0.45 μm-filtered and irradiated conditioned R10 with 45% fresh R10 and 25% FBS was used and feeding only occurred to offset media evaporation.

### RNA extraction, cDNA synthesis, and quantitative real-time polymerase chain reaction (RT-PCR) of lymphoblastoid cells

40-50 million cells were collected for each RNA extraction. Cells were pelleted for 5 minutes at 200 *x g,* washed in phosphate-buffered saline (PBS; Gibco), and resuspended in 1 ml TRIzol (Invitrogen). Cell lysis and phase separation with chloroform (Sigma) were performed as described from the TRIzol manufacturer protocol. Total RNA was isolated using the RNeasy Midi Kit (Qiagen) with in-column RNase-free DNase I (Qiagen) digestion. RNA was converted to cDNA using 4 μg of RNA per reaction, random hexamer primers, and the SuperScript III First-Strand Synthesis System (Invitrogen) according to manufacturer instructions. For quantitative RT-PCR, cDNA was amplified with primers SS_36F-SS_55R and SS_62F-SS_147R (exon-spanning when possible; sequences reported in Supplementary Table 2) and SsoFast EvaGreen Supermix (Bio-Rad) using the CFX Connect Real-Time PCR Detection System (Bio-Rad). Standard curves were employed routinely to quantify amplicons from each primer pair and assess expression of each respective gene in lymphoblastoid cells.

### Western blot analysis

25 million lymphoblastoid cells were collected for each protein extraction. Cells were pelleted for 5 minutes at 200 *x g,* washed in phosphate-buffered saline (PBS; Gibco), and resuspended in 200 μl Radioimmunoprecipitation assay buffer (RIPA; Sigma-Aldrich) with cOmplete protease inhibitors (Roche). After trituration and agitation for 30 minutes at 4°C, cellular debris was pelleted for 20 minutes at 17,000 *x g* and 6X Laemmli buffer (375 mM Tris-HCl, 9% SDS, 50% glycerol, 9% ß-mercaptoethanol, and 0.03% bromophenol blue [all Sigma]) was added to the supernatant to a final concentration of 1X. Samples were passed through a 27-gauge syringe 10 times and boiled at 95°C for 10 minutes. Samples and a positive control of human testis tissue lysate (Abcam, catalog number ab30257) were loaded into NuPAGE 4-12% Bis-Tris Protein gels (Novex) and run with NuPAGE MOPS SDS Running Buffer (Novex) in an XCell SureLock Mini-Cell Electrophoresis System (Novex) according to manufacturer instructions. Gels were transferred to 0.2 μm nitrocellulose membranes (Bio-Rad) using NuPAGE Transfer Buffer and an XCell II Blot Module (Invitrogen) overnight at 100 mA. Membranes were probed with primary antibodies anti-SYCP2 (EMD-Millipore, catalog number ABE2622) and anti-GAPDH (Cell Signaling Technology, catalog number 5174) and secondary antibody IRDye 800CW Donkey anti-Rabbit IgG (LI-COR). Membranes were visualized using the Odyssey Fc Imaging System (LICOR) and signal intensities were quantified in ImageJ (Version 1.46).

### Identification and evaluation of variable regions in *SYCP2* and *GAPDH*

*SYCP2* and *GAPDH* exons were compared against the database of Single Nucleotide Polymorphisms (dbSNP) to identify variable nucleotides^80^. To assess heterozygosity in DGAP230, genomic DNA of DGAP230 lymphoblastoid cells that had been pelleted and washed with phosphate-buffered saline (PBS, Gibco) was extracted using the DNeasy Blood and Tissue Kit (Qiagen). PCR was performed with LongAmp Taq 2X Master Mix (New England Biolabs, NEB) using customized primers (Integrated DNA Technologies). After amplification confirmation with agarose gel electrophoresis, and purification with the QIAquick PCR Purification Kit (Qiagen), Sanger sequencing reactions of PCR products were carried out with an ABI3730xl DNA analyzer. Chromatograms were aligned and multiple single nucleotide variants were called using Geneious (Version 7.0, Biomatters). The target exonic regions were selected based upon presence of multiple single nucleotide variants (amplified and Sanger sequenced with primers SS_174F-SS_175R and SS_192F-SS_193R from Supplementary Table 2). DGAP230 lymphoblastoid cell cDNA was acquired as described above under “RNA extraction, cDNA synthesis, and quantitative real-time polymerase chain reaction (RT-PCR) of lymphoblastoid cells” and assessed by the same method as the genomic DNA (amplified and Sanger sequenced with primers SS_175R-SS_193R as reported in Supplementary Table 2).

### 3C-PCR

3C-PCR was performed as previously described^26^. In brief, standard 3C libraries were generated^27,81–84^ and subjected to rearrangement-spanning nested PCR and subsequent Sanger sequencing using primers SS_216R-SS_227R (Supplementary Table 2).

### Circular Chromatin Conformation Capture sequencing (4C-seq) and analysis

4C-seq was adapted from previously described protocols with the following specifications^82–84^. Ten million cell aliquots of lymphoblastoid cells were crosslinked by adding formaldehyde (MilliporeSigma) to a final concentration of 2% for 10 minutes before the reaction was quenched with glycine (Promega) at a final concentration of 125 mM. After lysis, chromatin was further released by douncing with a 21.5-gauge needle and digested using 1500 U *HindIII* (New England Biolabs) per reaction at 37°C overnight. After enzyme inactivation, ligation, reverse crosslinking, DNA isolation, and purification, DNA was subjected to a second digestion by 50 U Csp6I (Thermo Fisher Scientific) per reaction at 37°C overnight. Enzyme inactivation, ligation, DNA isolation and purification were next performed to create the 4C libraries. Each library was amplified by inverse PCR with primers containing multiplexed overhangs (SS_198F-SS_212R as reported in Supplementary Table 2) using 50 ng of 4C template per reaction and purified by Qiaquick PCR purification kit (Qiagen). Samples were submitted to The Biopolymers Facility in the Department of Genetics at Harvard Medical School (Boston, MA, USA) for quality control on an Agilent 2200 TapeStation D1000 HS ScreenTape and by SYBR qPCR assay. Samples were spiked with 30% PhiX prior to sequencing on an Ilumina HiSeq 2500 sequencer using the HiSeq Single-Read Rapid Cluster Kit v2 (Illumina) for 100 cycles. At least 14 million pass-filter reads were acquired per sample.

4C-seq datasets were imported into the Galaxy-BioTeam Appliance (https://gbsc-galaxy.stanford.edu) and demultiplexed using Barcode Splitter (Galaxy Version 1.0.0)^85^. Reads were aligned to GRCh37/hg19 and a custom-made derivative chromosome 20 (der(20)) hg19 reference genome using Bowtie2 (Galaxy Version 0.2). The der(20) was made using a Perl script that parsed through chromosomes 20 and 22 and concatenated parts of the chromosomes at the resolved breakpoints (“code availability” section). Bam files were then analyzed using FourCSeq (Version 1.16.0) with normal parameters established by the FourCSeq protocol with the exception of using the der(20) for some analyses^31^. The interactions between fragments were then mapped to a circular plot using Circos (Version circos-0.69-6), by applying the relative coordinates of each fragment within the respective chromosome^86^. P-adjusted significance values were used to superimpose a colored heatmap onto the plot. These were then transformed into linear plots using Photoshop (Version CC 2015) by converting the image from Rectangular mode to Polar mode.

### CRISPR/Cas9-mediated genome editing in lymphoblastoid cells

CRISPR/Cas9-mediated deletion of chromosome 22 putative enhancers in the DGAP230 LCL was performed as previously described with the following parameters^34^. A Cas9-expressing stable DGAP230 cell line was developed by transduction with lentiCas9-Blast lentivirus, selection in blasticidin S HCl (Gibco), and validation by Western blot with primary antibodies anti-Cas9 (Abcam, catalog number 7A9-3A3) and anti-GAPDH (Cell Signaling Technology, catalog number 5174) and secondary antibodies IRDye 680RD Donkey anti-Mouse IgG and IRDye 800CW Donkey anti-Rabbit IgG (both LI-COR). This cell line was transduced with lentivirus derived from lentiviral vectors containing custom-made dual targeting sgRNAs that flank the putative enhancer region as well as a lentiGuide-Puro empty vector control. The dual targeting sgRNA vector was constructed and validated with primers SS_278F-SS_289F (Supplementary Table 2) and the previously described H1 bridge dsDNA block^34^. Selection of infected cells in puromycin (Gibco) yielded heterogeneous cultures, which were then either grown for one to four months or sorted into single cells in 96-well cell culture plates (Corning) using a MoFlo *Astrios* EQ cell sorter with a 100 μm nozzle at the Harvard Medical School Division of Immunology Flow Cytometry Core (Boston, MA, USA). Single cell clones were PCR-validated for homozygous deletions after genomic DNA extraction with the DNeasy Blood and Tissue Kit (Qiagen) by using LongAmp Taq 2X Master Mix (NEB) and primers SS_304F-SS_309R (Supplementary Table 2). To determine the relative amount of putative enhancer deletion in heterogeneous cultures derived from the dual targeting sgRNA lentivirus transduction and grown for different lengths of time, quantitative RT-PCR of 12 ng genomic DNA was performed with primers SS_308F-SS_309R and SS_52F-SS_53R (Supplementary Table 2) and SsoFast EvaGreen Supermix (Bio-Rad) using the CFX Connect Real-Time PCR Detection System (Bio-Rad). Standard curves were employed routinely to quantify amplicons from each primer pair.

### Construction of yeast strains

All strains used in this study (Supplementary Table 1) are isogenic to BR1919-8B^87^. Strains were created by standard genetic crosses and transformation procedures. For the development of strains AM3762 and AM4282, a *TRP1:P_GAL1_* promoter cassette was amplified from *pFA6a-TRP1-P_GAL1_*^88^ using Velocity polymerase (Bioline) and primers AJM1741-AJM1742 (Supplementary Table 2).

### Cytological analysis and imaging

Induction, sporulation, chromosome spreading, immunostaining, imaging and analysis were performed on diploid strains as previously described^89,90^ using the following parameters. Overnight cultures of YAM2592 (wild type) and AM3762 (*P_GAL1_-RED1*) were resuspended in 2% potassium acetate and split into two cultures per strain. Immediately after resuspension, one culture per strain was induced with 2 μm β-estradiol (Sigma E2257, prepared in DMSO) and the second received the corresponding volume of DMSO as an uninduced control. Cells were collected at 26 hours post sporulation and induction for chromosome spreading. The following primary antibodies were used for immunostaining at a 1:100 dilution: rabbit anti-Red1 (kind gift of G.S. Roeder^91^) and affinity-purified rabbit anti-Zip1 (raised at YenZym Antibodies, LLC, against a C-terminal fragment of Zip1^92^). The secondary antibody, donkey anti-rabbit conjugated to Alexa Fluor 488 (Abcam), was used at a 1:200 dilution. Imaging was carried out for eight different chromosome spreads derived from two distinct cultures per condition using a Deltavision RT imaging system (Applied Precision) adapted to an Olympus (IX71) microscope. Zip1 lengths were measured using the Softworx Measure Distance Tool.

### RNA extraction, cDNA synthesis, and quantitative RT-PCR of *S. cerevisiae*

Induction and sporulation were performed on diploid strains as previously described^89,90^. Overnight cultures of YAM2592 (wild type), AM4063 (wild type), AM3762 (*P_GALI_-RED1*), AM4282 (*P_GAL1_-RED1*), AM4283 (Δ*zip1*), AM4284 (Δ*red1*) and AM4286 (Δ*red1*) were resuspended in 2% potassium acetate and split into two cultures per strain. Immediately after resuspension, one culture per strain was induced with 2 μm β-estradiol (Sigma E2257, prepared in DMSO) and the second received the corresponding volume of DMSO as an uninduced control. Cells from 5 ml cultures were collected at 0, 6, and 26 hours post sporulation and induction and resuspended in 500 μl TRIzol (Invitrogen). RNA extraction, cDNA synthesis, and quantitative RT-PCR in S. *cerevisiae* were performed as described in the corresponding materials and methods section for LCLs with the exception of the following differences. Cells were disrupted by vortex with 0.5 mm glass beads (BioSpec Products) for 30 minutes at 4°C before phase separation by MaXtract High Density Tubes (Qiagen). Total RNA was isolated using the RNeasy Mini Kit (Qiagen) and RNA was converted to cDNA using 60 ng of RNA per reaction, oligo(dT) primers, and the SuperScript III First-Strand Synthesis System (Invitrogen). For quantitative RT-PCR, cDNA was amplified with primers SS_326F-SS_341R (sequences are reported in Supplementary Table 2). Standard curves were employed routinely to quantify amplicons from each primer pair and assess expression of each respective gene in S. *cerevisiae* cells.

### Münster male infertility cohort recruitment

Males attended the Centre of Reproductive Medicine and Andrology (CeRA) of the University Hospital Münster (UKM) to assess their fertility and were diagnosed by means of semen analysis performed according to the guidelines of the World Health Organization (WHO)^43^. After routine clinical diagnostics, males with known causes for their infertility including malignant disease, exposure to chemotherapy or radiation, structural and numerical chromosomal aberrations, and Y chromosome microdeletions were excluded from the study. All participants who gave written informed consent for evaluation of their clinical data and genetic analysis of their DNA samples were included in the MERGE (Male Reproductive Genomics) study of the Institute of Human Genetics, University of Münster. The study was approved by the Ethics Committee of the State Medical Board and the Medical Faculty in Münster (Kennzeichen 2010-578-f-S).

### Exome sequencing and analysis

Exome sequencing (ES) was performed in 627 patients with diverse infertility phenotypes from the MERGE study to identify possible deleterious sequence variants which might be causal for male infertility. Genomic DNA was isolated using standard procedures as previously described^64^. Samples were prepared, enriched, and indexed for ES according to the manufacturer’s protocol for SureSelect^QXT^ Target Enrichment for Illumina Multiplexed Sequencing Featuring Transposase-Based Library Prep Technology (Agilent). For multiplexed sequencing, libraries were index-tagged using appropriate pairs of index primers. To capture libraries, SureSelect^XT^ Human All Exon Kits (v4, v5 and v6) were used. Quantity and quality of libraries were determined with an Agilent TapeStation 2200 and the final concentration was adjusted to 1.6 pM. Sequencing was performed on the Illumina HiScan®SQ System, the Illumina NextSeq®500 System or the Illumina HiSeqX® System using the TruSeq SBS Kit v3 – HS (200 cycles), the NextSeq 500 V2 High-Output Kit (300 cycles) or the HiSeq Rapid SBS Kit V2 (300 cycles) respectively.

Reads were trimmed with Cutadapt v1.15^93^ and aligned to GRCh37.p13 using BWA-MEM v0.7.17^94^. Base quality recalibration and variant calling were performed using the GATK toolkit v3.8^95^ with haplotype caller according to their best practice recommendations. Resulting variants were annotated with Ensembl Variant Effect Predictor^96^.

Before evaluation of variants in *SYCP2,* participants with likely pathogenic or pathogenic variants in *TEX11, NR5A1,* and *DMRT1* were excluded from the study^64,65,67^. Rare variants in *SYCP2* were selected by a minor allele frequency (MAF) ≤1% in the gnomAD browser (http://gnomad.broadinstitute.org)^42^ and assessed for functional consequences at the protein level.

### Confirmatory Sanger sequencing

Identified variants in *SYCP2* were verified by Sanger sequencing according to standard procedures using primers SS_369F-SS_372R and SS_375F-SS_376R (Supplementary Table 2) and a 3730 DNA Analyzer (Applied Biosystems)^64^. When parental DNA was available, segregation was analyzed within the family using Sanger sequencing. Sequence analysis and visualization of the chromatograms was performed with CodonCode Aligner software (Version 8.0.1).

### Testicular biopsy histopathology

Testicular biopsies were fixed overnight in Bouin’s solution, washed with 70% ethanol, and embedded in paraffin. Subsequently, 5 μm sections were stained with Periodic acid-Schiff (PAS) and hematoxylin according to previously published protocols^97^. Slides were evaluated and documented using an Axioskop microscope (Zeiss, Oberkochen, Germany).

### Code availability

The GetDerivative.pl script used to create the der(20) and der(22) may be found at https://github.com/mnnshreya/Morton-DGAP230.

### Statistical analyses

For quantitative RT-PCR experiments, cDNA for three DGAP230 LCL replicates and three distinct age- and sex-matched control LCLs were each assessed in technical duplicates, normalized to *GAPDH* expression, and evaluated using an unpaired two-tailed t-test (Excel). For Western blot analysis, signal intensities of SYCP2 relative to GAPDH for three DGAP230 LCL replicates and three distinct age- and sex-matched control LCLs were assessed and evaluated using an unpaired one-tailed t-test (Excel). Statistical significance for RNA and protein analyses were determined by a p-value of p<0.05 and figure error bars show standard error of the mean. For identifying and assessing variable regions in DGAP230 LCLs, Sanger sequencing was performed on at least three different preparations of gDNA and/or cDNA. 3C-PCR and 4C-seq were each performed in triplicate, creating both 3C and 4C libraries from three independent cultures for the DGAP230 LCL and three different control LCLs. For differential analysis of 4C-seq datasets using FourCSeq (Version 1.16.0)^31^, read-counts were variance-stabilized and fit with a monotonic decay that was used to calculate z-scores. Z-scores were converted to p-values using a normal cumulative distribution curve and adjusted for multiple testing by the Benjamini-Hochberg method. A differential analysis was then conducted and significant interactions were detected by a p-adjusted value of p_adj_<0.01. For quantitative RT-PCR of the putative enhancer deletion, genomic DNA for two biological replicates of DGAP230 LCLs transduced with the dual targeting sgRNA lentivirus and grown for one month, 15 biological replicates of DGAP230 LCLs transduced with the dual sgRNA lentivirus and each grown for four months, and two biological replicates of DGAP230 LCLs transduced with the lentiGuide-Puro empty vector control were each assessed in technical triplicates, normalized to an intronic region of genomic DNA, and evaluated using a heteroscedastic one-tailed t-test for one versus four months and homoscedastic one-tailed t-test for dual targeting sgRNA lentivirus versus empty vector control (Excel). Statistical significance for quantitative RT-PCR analyses on genomic DNA was determined by a p-value of p<0.05 and figure error bars show standard error of the mean. For yeast cytology, at least 50 surface-spread meiotic nuclei were assessed per condition using eight different chromosome spreads derived from two distinct cultures. Graphpad Prism7 software was used for scatterplot generation of Zip1 filament length (with error bars indicating standard error of the mean) and statistical significance was determined by a Mann-Whitney U test, using VassarStats Concepts and Applications of Inferential Statistics (http://vassarstats.net/utest.html). Yeast qPCR experiments were performed in triplicate in two different strain backgrounds (N=6), normalized to *ACT1* expression, and evaluated using an unpaired one-tailed t-test for *RED1* and an unpaired two-tailed t-test for *ZIP1* (Excel). Statistical significance for RNA analyses were determined by a p-value of p<0.05 and figure error bars show standard error of the mean.

## Acknowledgements

This study was supported by the Eunice Kennedy Shriver National Institute of Child Health and Human Development (F31HD090780-01 to SLPS), the National Institute of General Medical Sciences (P01GM061354 to CCM, R15GM104827 and R15GM116109 to AJM), and the National Science Foundation (DGE1144152 to SLPS). The exome sequencing study was funded by the DFG Clinical Research Unit “Male Germ Cells: from Genes to Function” (CRU 326, DFG TU298/5-1). SJ is supported by the Leukemia & Lymphoma Society Career Development Program. DGAP sequencing reactions were carried out with an ABI3730xl DNA analyzer at the DNA Resource Core of the Dana-Farber/Harvard Cancer Center (funded in part by NCI Cancer Center support grant 2P30CA006516-48). Any opinion, findings, and conclusions or recommendations expressed in this material are those of the authors and do not necessarily reflect the views of the funding agencies.

## Author Contributions

S.L.P.S., F.T., A.J.M., and C.C.M. designed the study. S.L.P.S. performed all cellular, molecular, and genomic experiments, with the exception of jumping library genome sequencing, which was completed by C.H. and M.E.T., and exome sequencing and confirmatory Sanger sequencing of Münster subjects, which was completed by C.F. S.M. performed computational analyses for chromatin conformation capture sequencing and assisted with the CRISPR/Cas9-mediated deletion experiment. T.K., E.W., and S.K. ascertained and enrolled subjects and provided phenotypic information. S.K. also performed the testicular biopsy for participant M1401. S.J. provided guidance, protocols, and reagents for the CRISPR/Cas9-mediated deletion experiment. F.T. analyzed exome sequencing data from the Münster male infertility cohort and identified *SYCP2* frameshift variants. A.J.M. constructed yeast strains and provided guidance, protocols, and reagents for all yeast experiments. S.L.P.S. managed the project, initiated collaborations and wrote the manuscript with assistance from S.M., who wrote the methods and results sections for computational analysis of 4C-seq, C.F., who wrote the methods and results sections for the Münster male infertility cohort findings, and C.C.M., who edited the manuscript. The manuscript was approved by all authors.

## Materials and Correspondence

DGAP correspondence and requests for materials should be addressed to C.C.M. Yeast correspondence and requests for strains and AJM primers should be addressed to A.J.M. Correspondence regarding exome sequencing data and analyses should be addressed to F.T.

