## Supplementary information for "*SYCP2* translocation-mediated dysregulation and frameshift variants cause human male infertility"

### SUPPLEMENTARY RESULTS:

#### CRISPR/Cas9-mediated deletion of putative adopted enhancers

In order to implicate the genomic region identified by 4C-seq as the adopted enhancer for *SYCP2* dysregulation, we sought to determine causality by deleting the putative enhancer region using CRISPR/Cas9 and assessing the subsequent impact on *SYCP2* expression (Supplementary Fig. 1a)[1]. We established a Cas9-expressing stable DGAP230 LCL (Supplementary Fig. 1b) which were subsequently transduced with lentivirus derived from lentiviral vectors either containing dual targeting sgRNAs that flank the putative enhancer region or an empty vector control. Selection of dual targeting sgRNA-infected cells yielded a heterogeneous culture that demonstrates successful deletion of the target region (Supplementary Fig. 1c). Isogenic clones grown from single cell sorting revealed that only cells with biallelic intact enhancer regions survive (Supplementary Fig. 1d). To test whether the putative enhancer region is important for cell viability, dual targeting sgRNA-infected heterogeneous cultures were grown for variable lengths of time and quantitative RT-PCR of extracted genomic DNA was assessed for relative amounts of the targeted deletion. While the targeted deletion is enriched in dual targeting sgRNA-infected heterogeneous cells cultured for one month relative to the background signal from empty vector control-infected cultures, cells with the targeted deletion are selected out of the dual targeting sgRNA-infected heterogeneous population by four months (Supplementary Fig. 1e). These findings suggest that the putative enhancer region targeted by CRISPR/Cas9 is important for cell growth or viability. Due to the hypothesized biological significance of this region, we were unable to generate DGAP230 LCLs with a deleted putative enhancer region in order to study further its role in *SYCP2* overexpression.

**SUPPLEMENTARY FIGURES:**

**Supplementary Figure 1**

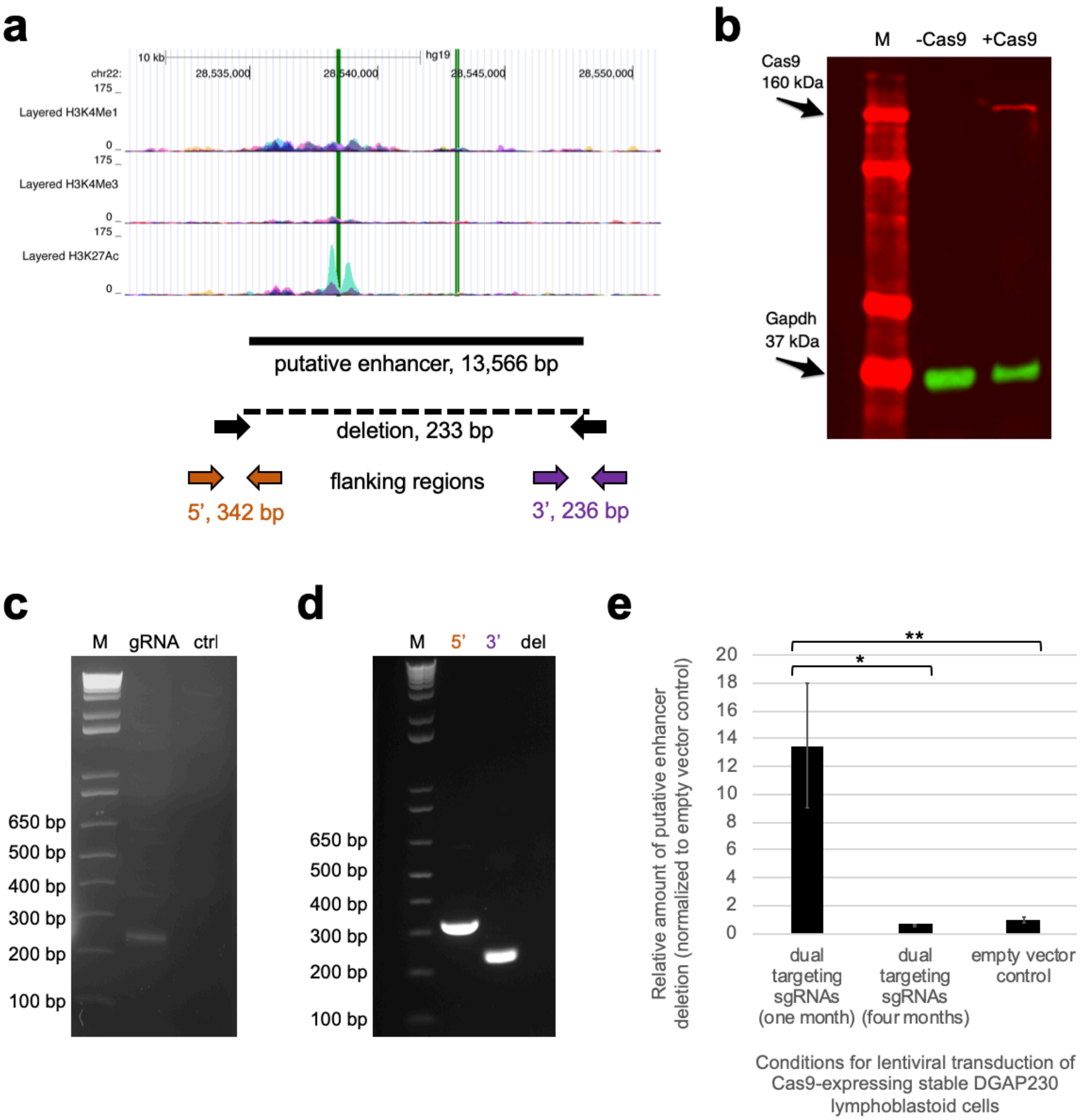


**Supplementary Figure 1: CRISPR/Cas9-mediated deletion of putative adopted enhancers.** **a,** The targeted deletion (black solid line) is mapped to the significant DNA fragment identified by 4C-seq. Tracks indicating enhancer activity from seven different cell lines are displayed in the UCSC genome browser and green bars highlight two regions of DNaseI hypersensitivity in fetal testis tissue according to ENCODE[2-4]. The dotted line and black arrows below the targeted deletion represent the predicted amplicon size by PCR of a deleted substrate. Orange and purple arrows represent the predicted PCR amplicon size of the 5' and 3' flanking regions, respectively.

**b,** A Cas9-expressing stable LCL was established for DGAP230, as indicated by Cas9 protein expression in a Western blot. M = marker; -Cas9 = uninfected DGAP230 LCL; +Cas9 = Cas9-expressing stable DGAP230 LCL. **c,** PCR amplification across the deletion in genomic DNA from the dual targeting sgRNA-infected culture reveals some CRISPR/Cas9-deleted target regions in the heterogeneous lymphoblastoid cell population. M = 1 kb plus DNA ladder; gRNA = deletion assessment in the heterogeneous culture infected with the dual targeting sgRNA lentivirus; ctrl = deletion assessment in the culture infected by the empty vector control lentivirus. **d,** Example of PCR analysis for an isogenic LCL grown from a single cell after dual targeting sgRNA-infection of Cas9-expressing stable DGAP230 lymphoblastoid cells. The presence of amplicons for the 5' and 3' flanking regions and absence of an amplicon across the deletion are interpreted as no deletion of the targeted enhancer region. M = 1 kb plus DNA ladder; 5' = 5' flanking region; 3' = 3' flanking region; del = deletion targeted by the dual targeting sgRNA lentivirus. **e,** Quantitative RT-PCR of the putative enhancer deletion in DGAP230 LCLs transduced with the dual targeting sgRNA lentivirus and grown for one month compared to four months show a loss of modified cells over time. Two biological replicates for the one-month and control conditions and 15 biological replicates for the four-month condition were each tested in triplicate and grouped for graphing and statistical analyses. Results are normalized to the empty vector control value and graphed as mean ± standard error. Results were found to be statistically significant between the one-month and four-month groups by a heteroscedastic one-tailed t-test ($p<0.02$) and between the one-month and control conditions by a homoscedastic one-tailed t-test ($p<0.01$). * = $p<0.05$; ** = $p<0.01$.

**Supplementary Figure 2**

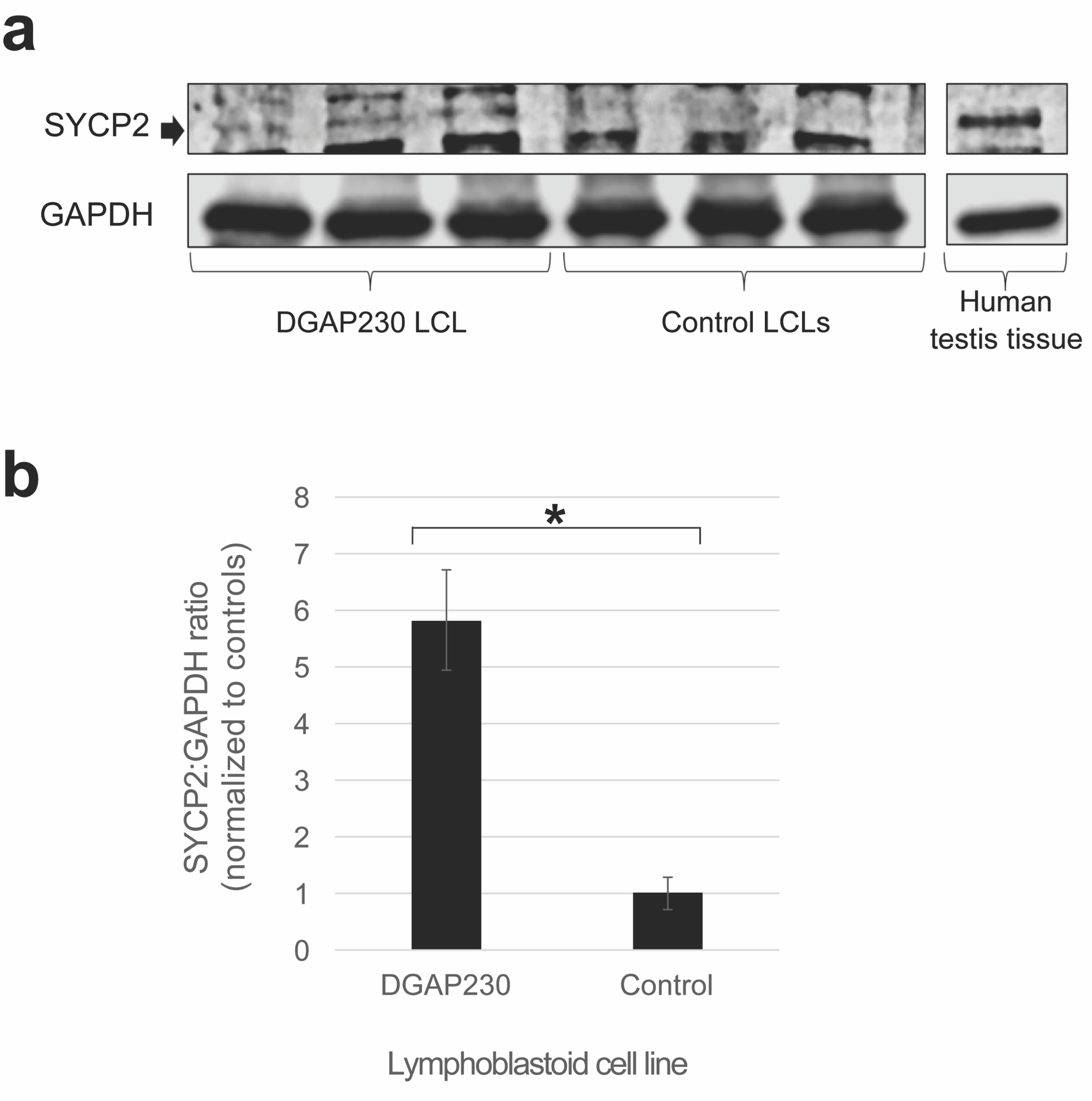


**Supplementary Figure 2: Western blot of synaptonemal complex protein 2 in DGAP230 and age- and sex-matched control lymphoblastoid cell lines. a,** The synaptonemal complex protein 2 (SYCP2) band, which has been confirmed by a human testis tissue lysate positive control, shows more abundant levels of expression in three biological replicates of DGAP230 lymphoblastoid cells (LCLs) compared to control LCLs from three different karyotypically normal fathers. **b,** Three technical replicates of each biological replicate were quantified in ImageJ (Version 1.46) and averaged. Biological replicates were then grouped into DGAP230 and control LCLs for graphing and statistical analysis. Results were found to be statistically significant by an unpaired one-tailed t-test ($N=3$; $p<0.0032$) and are graphed as mean ± standard error. * = $p<0.05$.

**Supplementary Figure 3**

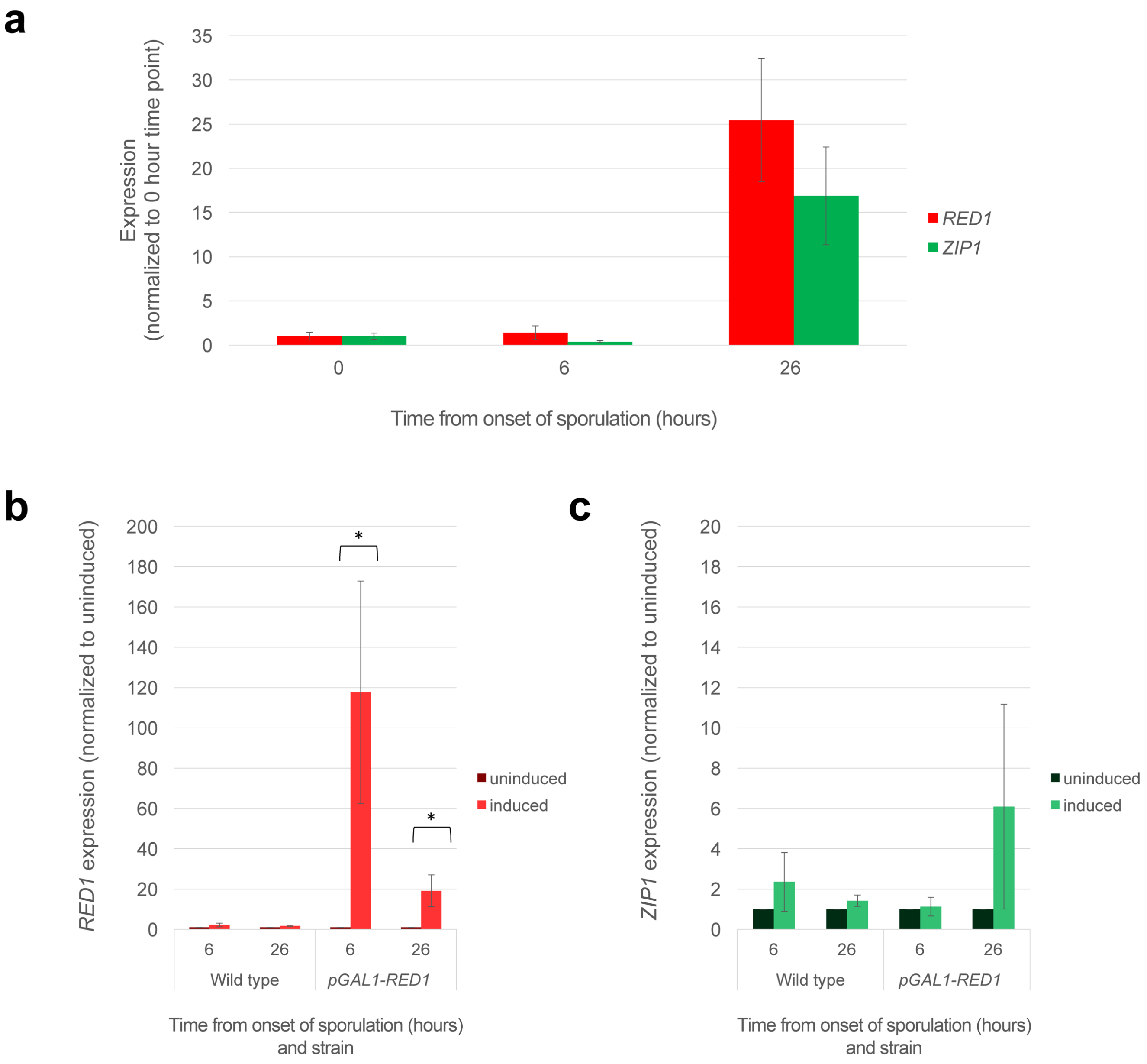


**Supplementary Figure 3: Control qPCR experiments for *RED1* and *ZIP1* transcription in sporulating *S. cerevisiae*. a,** Measurement of *RED1* and *ZIP1* transcript levels in wild type sporulating cells at various time points reveals gene induction between 6 and 26 hours after placement into sporulation medium. **b,** *RED1* is both constitutively expressed and overexpressed in the induced *pGAL1-RED1* strain background compared to uninduced *pGAL1-RED1*. These are statistically significant according to an unpaired one-tailed t-test ($p<0.0201$ and $p<0.026$, respectively), when tested in triplicate in two different strain backgrounds (N=6). * = $p<0.05$. **c,** *ZIP1* levels are not altered significantly with the induction of β-estradiol in either the wild type or *pGAL1-RED1* strains; differences were deemed not statistically significant according to an unpaired two-tailed t-test (all p-values between 0.327 and 0.687), when tested in triplicate in two different strain backgrounds (N=6). All results display mean ± standard error.

### SUPPLEMENTARY TABLES:

#### Supplementary Table 1. Yeast strains used in this study

| Strain Name | Alias | Genotype | Source | Notes |
|---|---|---|---|---|
| YAM1252 | wild type background | *lys2ΔNhe his4-260,519*<br>*lys2ΔNhe his4-260,519*<br><br>*leu2-3,112 MATα trp1-289*<br>*leu2-3,112 MAT**a** trp1-289*<br><br>*ura3-1 thr1-4 ade2-1*<br>*ura3-1 thr1-4 ade2-1* | Amy MacQueen | BR1919 background from[5] |
| YAM2592 and AM4063 | Ecm11-epitope tag heterozygote | YAM1252<br><br>*ECM11*<br>*ECM11-13MYC::kanMX4*<br><br>*ndt80Δ::LEU2*<br>*ndt80Δ::LEU2* | Amy MacQueen | For cytology and qPCR; at least one parent is different between the two strains (although identical in strain background and genotype) |
| AM3762 and AM4282 | $P_{GAL1}$-*RED1* | AM4063 *TRP1-*$P_{GAL1}$*-RED1*<br>*RED1*<br><br>*ura3::*$P_{GPD1}$*-GAL4(848).ER::URA3*<br>*URA3* | This study | For cytology and qPCR; at least one parent is different between the two strains (although identical in strain background and genotype) |
| AM4283 | Δ*zip1* | AM4063 *zip1Δ::URA3*<br>*zip1Δ::URA3* | Amy MacQueen | For qPCR negative control |
| AM4284 and AM4286 | Δ*red1* | AM4063 *red1Δ::HYG*<br>*red1Δ::HYG*<br><br>*ura3::PGPD1-GAL4(848).ER::URA3*<br>*URA3* | Amy MacQueen | For qPCR negative control; at least one parent is different between the two strains (although identical in strain background and genotype) |

**Supplementary Table 2. Primers used in this study**

| Name | Sequence 5'-3' | Target region | Description |
|---|---|---|---|
| SS_36F | TTCACCACCATGGAGAAGGC | *GAPDH* cDNA, human | For qPCR control with SS_37R; amplicon = 112 bp |
| SS_37R | TCTCATGGTTCACACCCATGAC | *GAPDH* cDNA, human | For qPCR control with SS_36F; amplicon = 112 bp |
| SS_40F | TTGGAAAAGGGACAGCCAAG | *SYCP2* cDNA, human | For qPCR with SS_41R; amplicon = 108 bp |
| SS_41R | GGTTGCTTTTCGTGGAAGTCTG | *SYCP2* cDNA, human | For qPCR with SS_40F; amplicon = 108 bp |
| SS_52F | CAGACTCCTGTGGTCAAGCA | *ATP2B1,* human | For qPCR with SS_53R; amplicon = 147 bp (control primer pair published in [6]) |
| SS_53R | TTCGTCAGTCAACCCCTTTC | *ATP2B1,* human | For qPCR with SS_52F; amplicon = 147 bp (control primer pair published in [6]) |
| SS_54F | TGACCACGTTTTCAGCTGTG | *CDH4* cDNA, human | For qPCR with SS_55R; amplicon = 103 bp |
| SS_55R | TTGGTGGCATTGATGTGCAG | *CDH4* cDNA, human | For qPCR with SS_54F; amplicon = 103 bp |
| SS_58F | GTTCGTGTCTCAGGTTCAGCCAG | upstream of DGAP230 rearrangement breakpoint on chr20 | For amplification with SS_59R of der(20) breakpoint; amplicon = 976 bp (also used for subsequent Sanger sequencing) |
| SS_59R | GCACAGTTTTGATCCTGTCTTGTGG | downstream of DGAP230 rearrangement breakpoint on chr22 | For amplification with SS_58F of der(20) breakpoint; amplicon = 976 bp (also used for subsequent Sanger sequencing) |
| SS_60F | GATAAGCCAATAACCACGACCTGAG | upstream of DGAP230 rearrangement breakpoint on chr22 | For amplification with SS_61R of der(22) breakpoint; amplicon = 832 bp (also used for subsequent Sanger sequencing) |
| SS_61R | GGATTAGGACAGGCAGGAGCAAG | downstream of DGAP230 rearrangement breakpoint on chr20 | For amplification with SS_60F of der(22) breakpoint; amplicon = 832 bp (also used for subsequent Sanger sequencing) |
| SS_62F | TGCCACAATGCACAGACAAC | *CDH26* cDNA, human | For qPCR with SS_63R; amplicon = 93 bp |
| SS_63R | TCTTCCAGCACATTGGCAAC | *CDH26* cDNA, human | For qPCR with SS_62F; amplicon = 93 bp |
| SS_70F | CCACCCAAACCTTAGAAAGCTG | *SNAP29* cDNA, human | For qPCR with SS_71R; amplicon = 100 bp |

| | | | |
|---|---|---|---|
| SS_71R | TCGAAGGTGTGGGTTC TTTG | *SNAP29* cDNA, human | For qPCR with SS_70F; amplicon = 100 bp |
| SS_76F | ACCAGCAACAGTATCG GGAAG | *SLC7A4* cDNA, human | For qPCR with SS_77R; amplicon = 85 bp |
| SS_77R | AGGCAGATGTTGAGGA CGATG | *SLC7A4* cDNA, human | For qPCR with SS_76F; amplicon = 85 bp |
| SS_92F | ACAGGTCAAGGTGTTC AACG | *PPP1R3D*, human | For qPCR with SS_93R; amplicon = 123 bp |
| SS_93R | AGGCAATGCAGGGTGA ACTC | *PPP1R3D*, human | For qPCR with SS_92F; amplicon = 123 bp |
| SS_94F | AGCACTACAGGAAAGA CAGTCC | *FAM217B* cDNA, human | For qPCR with SS_95R; amplicon = 120 bp |
| SS_95R | TGGGCCAGCATTCATA TTGC | *FAM217B* cDNA, human | For qPCR with SS_94F; amplicon = 120 bp |
| SS_98F | TTGTGAAATCGCGTGG ACTG | *C20orf197* cDNA, human | For qPCR with SS_99R; amplicon = 72 bp |
| SS_99R | TACCAACAGCTGCTCA GTGG | *C20orf197* cDNA, human | For qPCR with SS_98F; amplicon = 72 bp |
| SS_102F | AACATGATGCTGCCCA ACTG | *USP41* cDNA, human | For qPCR with SS_103R; amplicon = 138 bp |
| SS_103R | TGGCACAGTCAAGGCA AATC | *USP41* cDNA, human | For qPCR with SS_102F; amplicon = 138 bp |
| SS_106F | ATGGAGCATGCAGAGG GAAG | *ZNF74* cDNA, human | For qPCR with SS_107R; amplicon = 87 bp |
| SS_107R | TTGCAAATGCCCTGCT GTTG | *ZNF74* cDNA, human | For qPCR with SS_106F; amplicon = 87 bp |
| SS_110F | GCATGCCTGTAACCAT GTCAC | *SCARF2* cDNA, human | For qPCR with SS_111R; amplicon = 89 bp |
| SS_111R | TGCCATTGCTACACTTG GTC | *SCARF2* cDNA, human | For qPCR with SS_110F; amplicon = 89 bp |
| SS_114F | AACAACGATGCCGGAT ACAG | *KLHL22* cDNA, human | For qPCR with SS_115R; amplicon = 130 bp |
| SS_115R | TGTTGTCCAGCACAGC AATG | *KLHL22* cDNA, human | For qPCR with SS_114F; amplicon = 130 bp |
| SS_118F | AGCCATGTTTTCCTGAA GGC | *MED15* cDNA, human | For qPCR with SS_119R; amplicon = 137 bp |
| SS_119R | AGGCTCTGGAGTGCAT TCATAG | *MED15* cDNA, human | For qPCR with SS_118F; amplicon = 137 bp |
| SS_122F | AAGCGCCACTTCTCAG AAAC | *SERPIND1* cDNA, human | For qPCR with SS_123R; amplicon = 149 bp |
| SS_123R | TGAGCAGTTTCCCCTC CTTTC | *SERPIND1* cDNA, human | For qPCR with SS_122F; amplicon = 149 bp |
| SS_126F | CAGGTTTTGCATGTCCT CTTGG | *PI4KA* cDNA, human | For qPCR with SS_127R; amplicon = 122 bp |
| SS_127R | TTGCTGTAACGCCCAA ATGC | *PI4KA* cDNA, human | For qPCR with SS_126F; amplicon = 122 bp |
| SS_130F | AACCGCCGTTTTAAGAT CGG | *CRKL* cDNA, human | For qPCR with SS_131R; amplicon = 128 bp |

| | | | |
|---|---|---|---|
| SS_131R | ATTGGTGGGCTTGGAT ACCTG | *CRKL* cDNA, human | For qPCR with SS_130F; amplicon = 128 bp |
| SS_134F | TGAGGCCCAAGGAGTT TTTC | *AIFM3* cDNA, human | For qPCR with SS_135R; amplicon = 108 bp |
| SS_135R | TGAAGCCATCCTTGAA CACG | *AIFM3* cDNA, human | For qPCR with SS_134F; amplicon = 108 bp |
| SS_138F | AAGTTTGCAACTGGCC AGTG | *LZTR1* cDNA, human | For qPCR with SS_139R; amplicon = 109 bp |
| SS_139R | CAGCAAAGATCCACAG CTTGTC | *LZTR1* cDNA, human | For qPCR with SS_138F; amplicon = 109 bp |
| SS_142F | AAAGGACACAGTTACC CACCTG | *THAP7* cDNA, human | For qPCR with SS_143R; amplicon = 104 bp |
| SS_143R | GGTGGAGAAAATGGAG TTGTGG | *THAP7* cDNA, human | For qPCR with SS_142F; amplicon = 104 bp |
| SS_146F | TGACCAACTTCCTTGTG ACG | *P2RX6* cDNA, human | For qPCR with SS_147R; amplicon = 134 bp |
| SS_147R | TTTTACACCGTGGCTGT GTG | *P2RX6* cDNA, human | For qPCR with SS_146F; amplicon = 134 bp |
| SS_174F | TTGTGGACTCAACCCTT AGTCA | *SYCP2*, human | For PCR with SS_175R; amplicon = 2945 bp |
| SS_175R | GGACTTCCTCCCCCTT GTAA | *SYCP2*, human | For PCR with SS_174F; amplicon = 2945 bp (also used for subsequent Sanger sequencing) |
| SS_185F | GCTTGTGCGGATAGGT CAAT | *SYCP2* cDNA, human | For PCR with SS_186R; amplicon = 2404 bp |
| SS_186R | TGTTTCCAACAGTGTGC TGA | *SYCP2* cDNA, human | For PCR with SS_185F; amplicon = 2404 bp |
| SS_192F | TGAGCAGTCCGGTGTC ACTA | *GAPDH*, human | For PCR with SS_193R; amplicon = 619 bp for cDNA and 859 bp for gDNA (also used for subsequent Sanger sequencing) |
| SS_193R | TGACTCCGACCTTCAC CTTC | *GAPDH*, human | For PCR with SS_192F; amplicon = 619 bp for cDNA and 859 bp for gDNA |
| SS_198F | AATGATACGGCGACCA CCGAACACTCTTTCCCT ACACGACGCTCTTCCG ATCTTAAGGCGAGGGG TGCTGAGCTGGGTAC | *SYCP2* promoter, human | For 4C-seq inverse PCR with SS_212R; multiplexed for library "DGAP230_A2" using the barcode TAAGGCGA |
| SS_199F | AATGATACGGCGACCA CCGAACACTCTTTCCCT ACACGACGCTCTTCCG ATCTCGTACTAGGGGG TGCTGAGCTGGGTAC | *SYCP2* promoter, human | For 4C-seq inverse PCR with SS_212R; multiplexed for library "GM20184_B2" using the barcode CGTACTAG |

| | | | |
|---|---|---|---|
| SS_200F | AATGATACGGCGACCACCGAACACTCTTTCCCTACACGACGCTCTTCCGATCTAGGCAGAAGGGGTGCTGAGCTGGGTAC | *SYCP2* promoter, human | For 4C-seq inverse PCR with SS_212R; multiplexed for library "DGAP230_B1" using the barcode AGGCAGAA |
| SS_201F | AATGATACGGCGACCACCGAACACTCTTTCCCTACACGACGCTCTTCCGATCTTCCTGAGCGGGGTGCTGAGCTGGGTAC | *SYCP2* promoter, human | For 4C-seq inverse PCR with SS_212R; multiplexed for library "DGAP230_D3" using the barcode TCCTGAGC |
| SS_202F | AATGATACGGCGACCACCGAACACTCTTTCCCTACACGACGCTCTTCCGATCTGGACTCCTGGGGTGCTGAGCTGGGTAC | *SYCP2* promoter, human | For 4C-seq, inverse PCR with SS_212R; multiplexed for library "GM20188_A1" using the barcode GGACTCCT |
| SS_203F | AATGATACGGCGACCACCGAACACTCTTTCCCTACACGACGCTCTTCCGATCTTAGGCATGGGGGTGCTGAGCTGGGTAC | *SYCP2* promoter, human | For 4C-seq inverse PCR with SS_212R; multiplexed for library "DGAP278-02_A" using the barcode TAGGCATG |
| SS_212R | CAAGCAGAAGACGGCATACGACTTGGCCTTTCAAGCCGGC | *SYCP2* promoter, human | For 4C-seq inverse PCR with SS_198F-SS_203F |
| SS_216R | GAAGAGGAGCTTCTTAATGTACGC | *SYCP2*, human | For 3C-PCR second amplification with SS_220F; amplicon = 511 bp |
| SS_220F | ATTGCTTGAACCAGGAGGTG | chr22, proximal to rearrangement breakpoint | For 3C-PCR second amplification with SS_216R; amplicon = 511 bp (also used for subsequent Sanger sequencing) |
| SS_225F | TTTCTCTCCTGTTCCCAAGG | chr22, proximal to rearrangement breakpoint | For 3C-PCR first amplification with SS_227R; amplicon = 1843 bp |
| SS_227R | AGGTTGATCCTTGTTGAAATTGTT | *SYCP2*, human | For 3C-PCR first amplification with SS_225F; amplicon = 1843 bp |
| SS_278F | AAAGGACGAAACACCGCAGATACGAACAAAGAATCGGTTTTAGAGCTAGAAATAGCAAG | Plasmid lentiGuide-Puro and H1 bridge dsDNA block (underlined sequences); chr22 putative enhancer target region | For cloning of dual targeting sgRNAs into plasmid lentiGuide-Puro (Addgene plasmid #52963), used with SS_287R |
| SS_287R | TTCTAGCTCTAAAACCATTGTGGATCTGAGTGG | Plasmid lentiGuide-Puro and H1 bridge | For cloning of dual targeting sgRNAs into plasmid |

| | | | |
|---|---|---|---|
| | GAGGATCCAAGGTGTC TCATAC | dsDNA block (underlined sequences); chr22 putative enhancer target region | lentiGuide-Puro (Addgene plasmid #52963), used with SS_278F |
| SS_289F | GAGGGCCTATTTCCCA TGATT | Plasmid lentiGuide-Puro | For Sanger sequencing of sgRNAs inserted into plasmid lentiGuide-Puro (Addgene plasmid #52963) |
| SS_304F | CCTTTATCCCTAGGCA GCGT | chr22, 5' flanking region for chr22 putative enhancer deletion | For PCR with SS_305R; amplicon = 342 bp in cells with at least one intact chr22 putative enhancer |
| SS_305R | CAATCGCTTCAACTCCA CCC | chr22, 5' flanking region for chr22 putative enhancer deletion | For PCR with SS_304F; amplicon = 342 bp in cells with at least one intact chr22 putative enhancer |
| SS_306F | TCTTCTCCCTGGGCAT GAAC | chr22, 3' flanking region for chr22 putative enhancer deletion | For PCR with SS_307R; amplicon = 236 bp in cells with at least one intact chr22 putative enhancer |
| SS_307R | AGAGCCAGGATAAGAC TTGAGT | chr22, 3' flanking region for chr22 putative enhancer deletion | For PCR with SS_306F; amplicon = 236 bp in cells with at least one intact chr22 putative enhancer |
| SS_308F | TCCTTTATCCCTAGGCA GCG | chr22, across chr22 putative enhancer deletion | For PCR and qPCR with SS_309R; amplicon = 233 bp in cells with at least one deletion of the chr22 putative enhancer |
| SS_309R | TGACGATTCAGAGATTT GTGACT | chr22, across chr22 putative enhancer deletion | For PCR and qPCR with SS_308F; amplicon = 233 bp in cells with at least one deletion of the chr22 putative enhancer |
| SS_326F | TTGAGACTGGCATCGC AATG | *RED1, S. cerevisiae* | For qPCR with SS_327R; amplicon = 89 bp |
| SS_327R | TTTGGATTGAGGGACA CTGC | *RED1, S. cerevisiae* | For qPCR with SS_326F; amplicon = 89 bp |
| SS_332F | AACAAAAAGGAGGCGG ATGC | *ZIP1, S. cerevisiae* | For qPCR with SS_333R; amplicon = 96 bp |
| SS_333R | TTCGCTCAACTGACCT GAAC | *ZIP1, S. cerevisiae* | For qPCR with SS_332F; amplicon = 96 bp |
| SS_340F | TGGATTCTGAGGTTGC TGCTTTG | *ACT1* cDNA, *S. cerevisiae* | For qPCR with SS_341R; amplicon = 103 bp |
| SS_341R | ACGATAGATGGGAAGA CAGCAC | *ACT1* cDNA, *S. cerevisiae* | For qPCR with SS_340F; amplicon = 103 bp |

| | | | |
|---|---|---|---|
| AJM1741 | ATTTTTTAATCAGTGAGGACCACAAAGGGACAGCAAATACGGTGATAAGAGAATTCGAGCTCGTTTAAAC | *pFA6a-TRP1-*$P_{GAL1}$ (underlined sequence) and *RED1* promoter, *S. cerevisiae* | Forward primer to amplify *TRP1-*$P_{GAL1}$ and place it upstream of *RED1*. Used with AJM1742 |
| AJM1742 | AAGTCATTTTTCAGGCAAACACCAAAAATCTTTTTCTTCAAACCTTCCATTTTGAGATCCGGGTTTT | *pFA6a-TRP1-*$P_{GAL1}$ (underlined sequence) and *RED1* promoter, *S. cerevisiae* | Reverse primer to amplify *TRP1-*$P_{GAL1}$ and place it upstream of *RED1*. Used with AJM1741 |
| AJM1743 | ACGATTTCGCAGCAGGATCAGATGG | *RED1* promoter, *S. cerevisiae* | For genotyping of *TRP1-*$P_{GAL1}$*-RED1* |
| SS_369F | TGGGCCATGAGTAGACAAGTG | *SYCP2*, human | For PCR with SS_370R to amplify variant c.2793_2797del in exon 31; amplicon = 703 bp (also used for Sanger sequencing) |
| SS_370R | ACCTCTCTGGAAATAAGTTGTTTTGA | *SYCP2*, human | For PCR with SS_369F to amplify variant c.2793_2797del in exon 31; amplicon = 703 bp (also used for Sanger sequencing) |
| SS_371F | TCACTAGATTCAGACATCTGTTTTG | *SYCP2*, human | For PCR with SS_372R to amplify variant c.3067_3071del in exon 33; amplicon = 564 bp (also used for Sanger sequencing) |
| SS_372R | GATGATAACTGGAATGGGAAGAT | *SYCP2*, human | For PCR with SS_371F to amplify variant c.3067_3071del in exon 33; amplicon = 564 bp (also used for Sanger sequencing) |
| SS_375F | AGGCCAATCACTCTGCTTGG | *SYCP2*, human | For PCR with SS_376R to amplify variant c.2022_2025del in exon 24; amplicon = 399 bp (also used for Sanger sequencing) |
| SS_376R | GCAAGTACTTCGTCAGGAGACA | *SYCP2*, human | For PCR with SS_375F to amplify variant c.2022_2025del in exon 24; amplicon = 399 bp (also used for Sanger sequencing) |
